# Model-based decoupling of evoked and spontaneous neural activity in calcium imaging data

**DOI:** 10.1101/691261

**Authors:** Marcus A. Triplett, Zac Pujic, Biao Sun, Lilach Avitan, Geoffrey J. Goodhill

**Affiliations:** Queensland Brain Institute, The University of Queensland, St Lucia, QLD 4072, Australia; School of Mathematics and Physics, The University of Queensland, St Lucia, QLD 4072, Australia

## Abstract

The pattern of neural activity evoked by a stimulus can be substantially affected by ongoing spontaneous activity. Separating these two types of activity is particularly important for calcium imaging data given the slow temporal dynamics of calcium indicators. Here we present a statistical model that decouples stimulus-driven activity from low dimensional spontaneous activity in this case. The model identifies hidden factors giving rise to spontaneous activity while jointly estimating stimulus tuning properties that account for the confounding effects that these factors introduce. By applying our model to data from zebrafish optic tectum and mouse visual cortex, we obtain quantitative measurements of the extent that neurons in each case are driven by evoked activity, spontaneous activity, and their interaction. This broadly applicable model brings new insight into population-level neural activity in single trials without averaging away potentially important information encoded in spontaneous activity.

## Introduction

The nervous system constructs internal representations of its sensory environment through coordinated patterns of neural activity. Uncovering these representations from neural recordings is a central problem in systems neuroscience. Typically this task is approached by measuring the relationship between the parameters of a stimulus and the neural response following stimulus presentation. However, the pattern of neural activity evoked by a stimulus is highly variable, and can be different each time the stimulus is presented. An important source of this variability is ongoing spontaneous activity (SA) that does not appear to be driven by the stimulus [1]. In some cases this SA may simply be biophysical noise that should be averaged away, but in other cases it may represent salient features of brain function such as parallel encoding of non-sensory variables [2, 3], mechanisms for circuit development [4], or other internal-state factors that regulate sensory-guided behaviour [5]. Uncovering the interplay between stimulus-evoked activity (EA) and SA therefore requires the ability to reliably separate these two components. This is challenging, however, because the internal factors that give rise to SA are often unknown or cannot currently be directly measured.

This problem is particularly acute for calcium imaging data, a major source of our current understanding of the joint activity of large numbers of neurons. Neurons that express calcium indicators report activity at a high spatial resolution, but filter out high frequency spiking due to slow indicator binding kinetics and saturating calcium concentrations [6, 7]. These calcium levels in turn are only observed through subsampled fluorescence intensities that are subject to noise from the optical imaging system. Moreover, neurons can be recorded in large populations with many thousands of imaging frames, leading to very high dimensional data that can challenge traditional methods of neural data analysis [8].

Much research in recent years has focused on statistical methods for extracting hidden (or “latent”) structure from neural population data [9–12]. A key assumption in these methods is that neural population activity tends to possess a characteristic low dimensional structure, reflecting underlying constraints on how neurons can comodulate their activity [13]. Thus high dimensional neural data can often be well-described by a much smaller number of latent variables evolving through time. In this context, unobserved sources of SA are latent variables that can be inferred from data given the appropriate statistical tools. However, methods for identifying latent structure in calcium imaging data (see e.g. refs. [14–16]) are scarce compared to spike train data and none so far have sought to explicitly extract sources of SA hidden amidst population responses to sensory stimuli.

Here we develop a latent variable model for calcium imaging data that allows for a decomposition of single-trial neural activity into its evoked and spontaneous components. In our model, which we refer to as calcium imaging latent variable analysis (CILVA), patterns of SA are driven by hidden factors decoupled from the stimulus. By fitting the model to data, we identify the structure and temporal behaviour of these latent sources of SA, and simultaneously extract receptive fields that are not biased by the variability that these sources of SA introduce. To demonstrate the applicability of the model we analysed calcium imaging data from both the larval zebrafish optic tectum and mouse visual cortex. In both cases we identified sparsely active independent latent factors that targeted distinct sets of neurons. Besides revealing the statistical structure of SA, accounting for these factors produced sharper receptive field estimates, more refined retinotopic maps, and quantitative measurements of the presence and interaction of EA and SA.

## Results

### Low dimensional spontaneous activity proceeds throughout stimulus presentation

As a first example we considered two-photon calcium imaging data from the optic tectum of the developing larval zebrafish. Fish expressing the genetically encoded calcium indicator GCaMP6s were embedded in low melting point agarose while small dark spots were presented at systematically varying angles across the visual field. Onset of the visual stimulus evoked calcium transients in the tectum (Figure 1a) consistent with the topography of the retinotectal map. The presentation of a spot was followed by an interval of 19s without any stimulation, allowing calcium levels to return to baseline before the next stimulus. The optic tectum was often highly active during these inter-stimulus intervals (Figure 1b) and calcium transients sometimes occurred spontaneously just before stimulus onset, elevating the recorded fluorescence levels associated with that stimulus.

**Figure 1:**
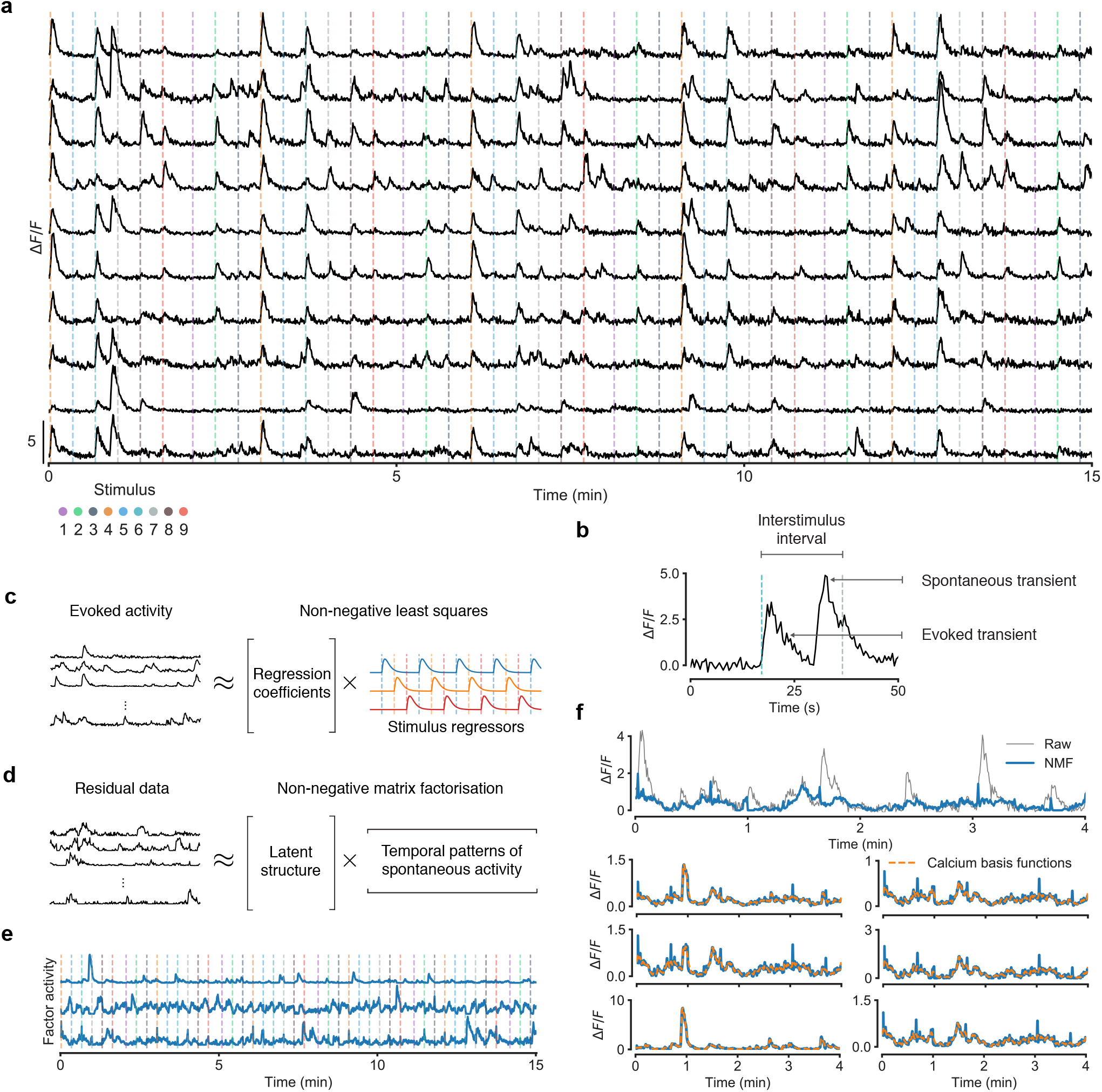
Two-photon calcium imaging and application of the residual NMF approach to the larval zebrafish optic tectum. **a**, Fluorescence traces from 10 example neurons. Dashed vertical lines indicate stimulus onset; colour represents azimuth angle of presented stimulus. **b**, Example fluorescence trace segment illustrating that spontaneous calcium transients can occur just before stimulus onset. **c**, A simple estimate of the stimulus-driven component of population data can be obtained by multiple regression of fluorescence traces onto stimulus regressors using non-negative least squares. **d**, After estimating the stimulus-driven component, low dimensional structure in the residual data can be estimated using non-negative matrix factorisation. **e**, Patterns of SA shared between groups of neurons found via NMF. For consistency with later results, we here applied NMF with three latent factors. Each row corresponds to the activity of one factor. **f**, Top: component of the raw fluorescence trace (black) considered to be SA by the residual NMF approach (blue). NMF often produces estimates with erratic and sudden changes in calcium levels that fail to respect the stereotypical structure of calcium activity. Bottom: additional examples of shared SA estimated from the residuals using NMF (blue). For comparison, the same estimates are shown when expressed in a basis of calcium impulse response functions located at each time point (orange, Methods). Deviations from the orange curve demonstrate atypical calcium behaviour. Samples were selected for illustration from among the 10 neurons best explained by the residual NMF approach.

Before adopting a model-based approach to the separation of spontaneous from evoked activity, we first considered using a combination of existing tools. We characterised the stimulus-driven component of the population activity by simply expressing each fluorescence trace as a linear combination of regressors that specify the basic shape and timescale of calcium activity. We defined a basis of stimulus regressors [17, 18] by convolving a calcium impulse response kernel with the presentation times of each stimulus and performed a multivariate linear regression of the population data onto this basis set using non-negative least squares (Figure 1c). Any highly structured activity in the residual data would likely be driven primarily by latent sources of SA not directly related to the onset of the presented stimuli. We then applied non-negative matrix factorisation [19] (NMF) to search for low dimensional structure in the residuals. NMF attempted to reconstruct the residual population data as the product of two matrices with non-negative entries: a matrix whose rows are timeseries that capture patterns of SA shared across groups of neurons, and a matrix whose columns describe how neurons are coupled to such timeseries (Figure 1d). The NMF description of SA identified low dimensional structure that proceeded throughout the recording, largely independent of the stimulus (Figure 1e).

While this residual NMF approach is efficient due to highly optimised computational routines, it is limited by two characteristics. First, because the models of stimulus processing and shared SA are not inferred jointly, receptive field estimates are biased towards higher values by spontaneous calcium transients that coincide with stimulus presentations. This in turn will introduce bias when applying NMF to the residual data, since some of the contribution of SA will have already been subtracted out at the receptive field estimation stage. Second, NMF has no model for the highly stereotyped structure of calcium transients, and therefore does not respect this structure in the components that it finds (Figure 1f).

### CILVA simultaneously captures evoked responses and shared spontaneous activity

To overcome these difficulties, we developed a generative statistical model that describes evoked and spontaneous activity simultaneously rather than sequentially (Figure 2a,b). The model specifies the observed fluorescence level *f_n_*(*t*) for neuron *n* at time *t* in terms of the underlying calcium concentration *c_n_*(*t*),

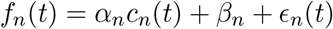

where the scalars *α_n_* and *β_n_* determine the scale and baseline of the fluorescence signal respectively, and *ϵ_n_*(*t*) represents Gaussian noise. Consistent with experimental data [6] and previous models for calcium imaging [7, 20], the calcium dynamics are assumed to be highly stereotyped and are defined by the convolution of a calcium impulse response kernel **k** with an activity intensity function ***λ***_*n*_ (characters in bold represent vectors or matrices)

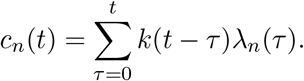

The kernel **k** is a difference-of-exponentials function (see Methods), which includes both rise and decay time constants. We found that including an explicit rise time was essential as GCaMP6s activity is slow relative to the sampling rate of many optical imaging systems.

**Figure 2:**
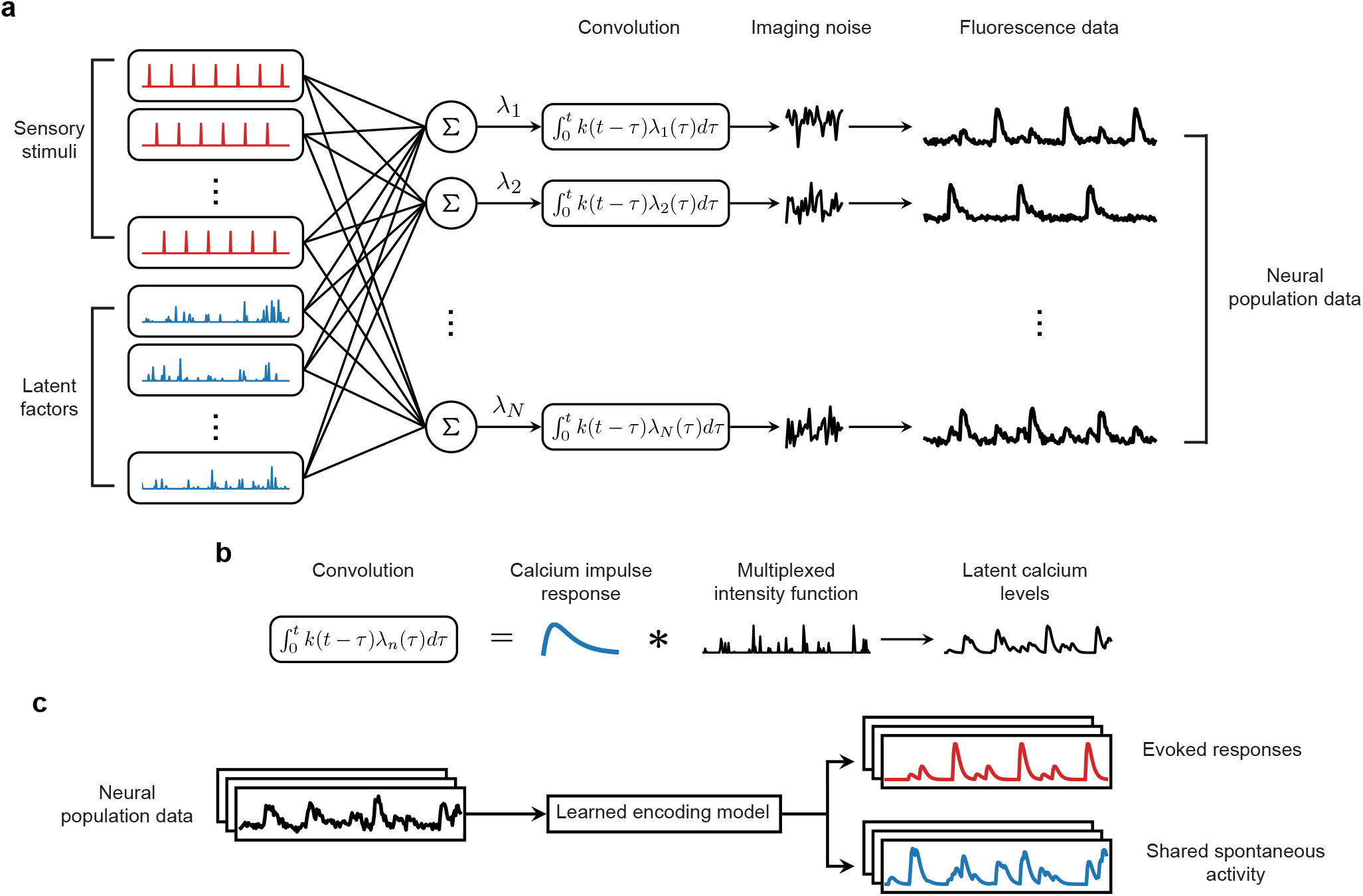
Overview of the CILVA approach for decoupling stimulus-evoked responses and latent sources of SA. **a**, Proposed generative architecture underlying multivariate calcium imaging data. Neurons are driven by sensory stimuli (red) and latent sources of SA (blue). These two sources are combined additively to define the underlying intensity of neural activity (***λ***_*n*_), before being convolved with a calcium kernel. Calcium levels are subsequently reported through noisy fluorescence intensities. **b**, Intensity functions ***λ***_*n*_ encoding stimuli and shared SA are convolved with a calcium impulse response kernel **k** to generate calcium levels. **c**, The learned encoding model provides a method for decoupling evoked responses from common patterns of SA.

The key ideas of the model are that (i) evoked responses will tend to be locked to the onset of the stimulus, (ii) evoked responses typically have a simple impulse response structure in calcium imaging data, and (iii) neural activity not attributable to evoked activity should be explained as far as possible by SA with a specific structure. Neural activity *λ_n_*(*t*) in each imaging frame *t* was thus assumed to be driven by the addition of two underlying sources: processing of the stimulus **s**(*t*) through a linear receptive field **w**_*n*_, and a small number of unobserved or “latent” sources of SA **x**(*t*),

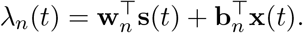

Here **x**(*t*) is the low dimensional latent state at time *t*, **b**_*n*_ is a vector describing how neuron *n* is affected by these factors, and ^⊤^ denotes the transpose operation. This SA model is inspired by the application of factor analysis methods to neural population data [9, 21], which posit that low dimensional structure arises from latent computations or brain states that concurrently affect subsets of neurons. However, the latent factors underlying spontaneous calcium transients in our model differ mathematically from classical factor analysis in that, due to non-negativity of the calcium levels, factor activity states and coupling between latent factors and neurons must be non-negative [22].

We fit the model by computing the maximum a *posteriori* estimate of the latent factor activity states. Because these activities were less constrained by the model compared to the time-locked evoked responses (and therefore likely to be more complex) we encouraged sparsity by placing a non-negative prior on the latent factors with high density near zero, and used a simple model selection procedure to estimate the sparsity penalty (see Methods).

### Fluorescence signals can be decomposed into their evoked and spontaneous components

The formulation of our model is convenient for analysing the separate contributions of evoked and spontaneous activity to the observed fluorescence levels (Figure 2c). The fitted model can be succinctly summarised by the equation

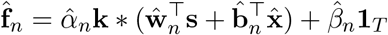

where 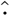 denotes an estimated variable, * denotes linear convolution, and **1**_*T*_ is a vector of ones with length equal to the number of imaging frames *T*. Here we have dropped explicit dependence on the calcium levels *c_n_*(*t*), which are deterministic given the other model parameters. The components of the signal driven purely by evoked or spontaneous activity can then be extracted from the convolution to give

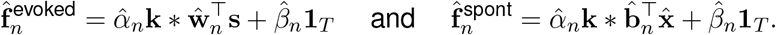

We first verified the model on simulated data with known ground truth, modelling the properties of the zebrafish and mouse data that we subsequently consider (Supplementary Tables 1 and 2, Supplementary Figures 1 and 2). We then applied the model to our calcium imaging data from the zebrafish optic tectum to decouple the evoked and spontaneous calcium transients (Figure 3). Overlaying these decoupled calcium traces onto the experimental data we found that they provided realistic descriptions of calcium activity (Figure 3a) and a close fit between the raw fluorescence trace and the model reconstruction (Figure 3b, Supplementary Figure 3). The time-locked responses to stimuli were well modelled by 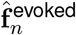, while low dimensional SA was identified by the projected latent factor activity 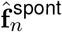 (Figure 3a, Supplementary Figure 4, Supplementary Videos 1 and 2). Neural activity in the residual data (i.e., after subtracting the model reconstruction from the raw fluorescence traces) arose primarily from spontaneous calcium transients that were independent of the latent sources of shared SA (and were therefore attributed to private variability [23], Supplementary Figure 5).

**Figure 3:**
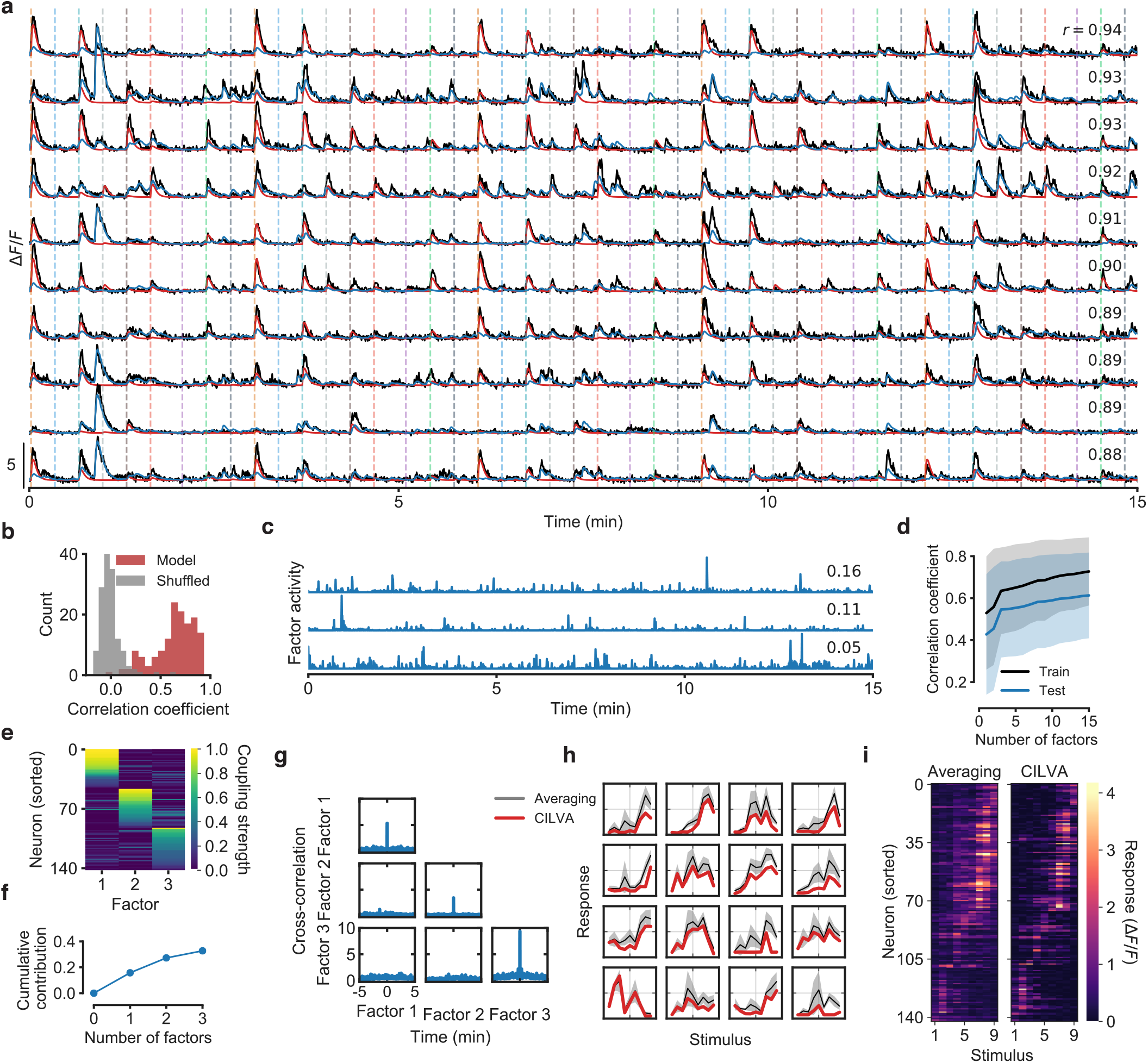
Fitted model components for the zebrafish shown in Figure 1. **a**, Results of fitting CILVA and decoupling EA (red) and shared SA (blue) in an experimental recording. Inset numbers denote the Pearson correlation coefficient between raw fluorescence trace and model fit. The 10 neurons with the highest correlations between data and model fit are shown. **b**, Distribution of correlation coefficients between data and model fits. Shuffled data obtained by cyclically permuting each trace by a random offset while preserving its temporal structure. **c**, Inferred latent factor timeseries. Inset numbers denote the factor contribution indices, defined as the mean reduction in correlation coefficient across the population following deletion of the corresponding factor. **d**, Cross-validated distributions of correlation coefficients for 1 to 15 latent factors. Shaded error bars indicate one standard deviation. For this fish adding additional latent sources of SA beyond three factors provides little improvement in the average correlation for both training and held-out test data. **e**, Estimated factor coupling matrix shows that latent factors target distinct, nonoverlapping sets of neurons. **f**, Cumulative factor contribution indices for 0 to 3 latent factors. **g**, Cross-correlograms show little interaction between latent factors and no long term structure in individual factor activity. **h**, Example estimated stimulus tuning curves (red). Tuning curves obtained by simply trial-averaging fluorescence levels over a window following stimulus onset (gray) provided for comparison (Methods). Shaded error bars denote one standard deviation. **i**, Retinotopic maps obtained by trial-averaging fluorescence levels over a window following stimulus onset (left) and by CILVA-estimated tuning curves (right). The CILVA map is more refined since the estimated stimulus filters already account for ongoing SA.

We fit the model with three latent factors, whose inferred activity timeseries were sparse (Figure 3c). Including additional latent factors beyond these resulted in better models of the SA of individual neurons or small subsets of neurons, but caused little improvement in the overall quality of model fit for this fish (Figure 3d). To understand the relative importance of each factor to the overall model fit, we defined a contribution index for a factor as the average reduction in the quality of model fit following its deletion (Methods). We found that each factor made a substantial contribution to the overall model fit by modulating shared SA across large groups of tectal neurons (Figure 3e,f).

The factor coupling matrix (defined by the vectors **b**_*n*_) reports how neurons are affected by the latent factors. There are several possibilities for how this matrix could be structured. First, if there is a minimal presence of structured SA the coupling matrix may exhibit no coherent organisation at all. Second, neurons could require the coordinated activity of several latent factors to explain their SA. This would be the case if, e.g., neurons participated in multiple recurrently connected circuits driven by noise [24, 25], and would result in factors modulating overlapping groups of neurons. Third, latent factors may each drive their own distinct sets of neurons, with little cross-talk between them. This could occur if, e.g., latent factors were encoding unrelated streams of motor or non-visual sensory information [2, 26]. In this larval zebrafish the estimated coupling matrix had a highly modular structure, with factors influencing largely non-overlapping sets of neurons (Figure 3e). Furthermore, the factor cross-correlograms (Figure 3g) showed no sign of dependence between factors, indicating that distinct sets of neurons were uniquely targeted by independent latent sources of SA.

Since our model fits receptive fields jointly with latent sources of SA, the estimated tuning curve for each neuron already accounts for ongoing SA that may have inflated its responses to stimuli. Indeed, if spontaneous calcium transients coincide with the presentation of a stimulus, one could expect tuning curves obtained by simply averaging the fluorescence levels over a small window following stimulus onset to be spuriously larger, and exhibit higher variance than if these events did not occur. We plotted the tuning curves estimated by CILVA against tuning curves obtained by averaging (see Methods) and found that they confirmed this intuition (Figure 3h). Moreover, sorting the neurons according to their preferred stimulus revealed a more refined retinotopic map when explicitly accounting for SA (Figure 3i). While our data showed a variety of tuning types, of note were neurons that were insensitive to visual stimuli (i.e., had relatively flat tuning curves), but that were highly active throughout the recording.

### Neurons are differentially driven by external stimuli and latent internal factors

To quantify the extent to which each neuron is driven by sensory stimuli versus shared SA, we derived an equation that expressed the variance of the reconstructed fluorescence levels in terms of three components: the variance attributable solely to EA, the variance attributable solely to shared SA, and the covariance (i.e., interaction) between EA and shared SA (Figure 4a, Methods). We defined a “drive ratio” to measure whether neurons were driven more by SA or EA (Methods). This revealed that across the population there was a continuous progression of responses, with some neurons being primarily driven by EA, some primarily by SA, and some by a mixture of both SA and EA (Figure 4b). To confirm that these effects were not artefacts of the model or the calcium indicator, we verified that the model does not overestimate the variance in the data (Figure 4c) and that there were interactions between EA and SA that were greater than expected by chance (Figure 4d). The distribution of drive ratios was largely bimodal (Figure 4e), indicating a preference to be dominated by either EA or SA rather than responding equally to both. There was no systematic relationship between quality of model fit and drive ratio (Figure 4f), and as expected, neurons strongly driven by sensory stimuli were weakly coupled to latent factors (Figure 4a,g).

**Figure 4:**
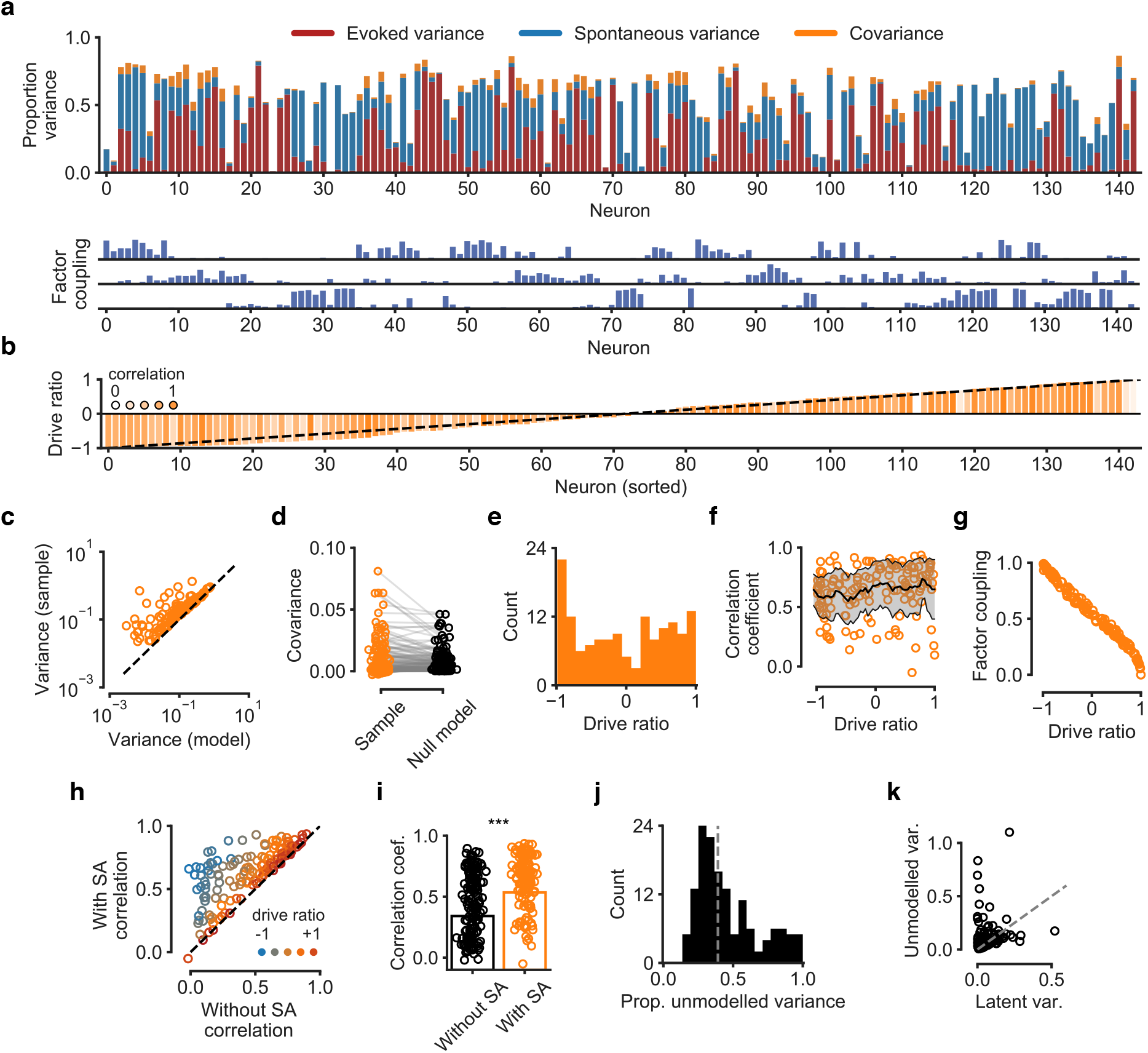
Analysis of the contribution of EA and SA to neural variability. All data for the same fish as in Figure 3. **a**, Top: composition of each neuron’s sample variance in terms of variance attributable solely to EA (red bars), solely to shared SA (blue bars), and their covariance (orange bars). Orange bars represent absolute values of covariances for ease of visualisation. Variance components are given as proportions of the total sample variance of the raw fluorescence signal var[**f**_*n*_] (corrected for imaging noise, see Methods). Neurons sorted roughly by their position on the anterior-posterior axis of the optic tectum. Bottom: coupling between neurons and latent sources of SA suggests neurons with strong coupling are weakly driven by sensory stimuli. Maximum bar height of one. **b**, Drive ratios vary gradually from −1 to +1, indicating a progressive shift in the composition of neural activity from purely spontaneous to purely evoked. For comparison, the dashed diagonal line represents a uniform distribution of drive ratios. **c**, Sample variance (corrected for imaging noise) vs variance of the statistical model indicates that the model does not overestimate variance. Each data point represents one neuron. **d**, Covariances between evoked and spontaneous traces estimated by the model (orange circles). Chance levels (black circles) are 95th percentiles of shuffled data obtained by cyclically permuting evoked traces by random offsets 1000 times while preserving temporal structure. Sample covariances exceeding chance levels cannot be attributed to the slow timescale of the calcium indicator. **e**, Distribution of drive ratios across the population of neurons. **f**, Correlation coefficients between experimental data and model fit as a function of drive ratio shows the model is not biased towards fitting spontaneous over evoked traces or vice versa. Black line represents the mean correlation coefficient in a small sliding window centred at the corresponding drive ratio; shaded region represents one standard deviation from the mean. **g**, Overall factor coupling (defined as the Euclidean norm ║**b**_*n*_║_2_ for each neuron *n*) as a function of drive ratio. Neurons strongly driven by sensory stimuli are typically weakly affected by latent sources of SA, and vice-versa. **h**, Correlation coefficient between raw fluorescence trace and model fit with and without incorporation of SA. Neurons with strongly negative drive ratios show marked improvement in the quality of model fit. **i**, Statistically significant improvement in the average correlation coefficient between experimental data and model fits after incorporating latent sources of SA (*p* < 0.001, Wilcoxon signed-rank test). **j**, Distribution of unmodelled variance, as a proportion of (corrected) sample variance. Dashed line indicates a median of 0.39. **k**, Variance attributable to latent sources of SA (latent variance) vs unmodelled variance identifies a population of neurons whose variance arises primarily from unobserved common input rather than private variability; i.e., neurons below the unity line (dashed).

We next quantified the improvement that resulted from incorporating SA into the model (Figure 4h,i). Without SA, the model for each neuron consists of a simple linear filter convolved with a calcium kernel. This well-describes neurons that possess high drive ratios (i.e., whose variances are dominated by EA) and thus these neurons show little improvement in how well the statistical model fits their fluorescence levels with the incorporation of SA (Figure 4h, neurons along the diagonal). In contrast many neurons are poorly fit by a model that incorporates only stimulus responses, and show substantial improvement when shared sources of SA are taken into account (Figure 4h, neurons above the diagonal), leading to a significant increase in the average correlation between fluorescence traces and model fits (Figure 4i).

Our model provides a simple approach to estimating the relative presence of shared vs private variability. Variance that is not captured by the model (Figure 4j) can be attributed to either private neural variability or residual imaging noise. Unmodelled variance thus represents an upper bound on private variability. We plotted the variance attributable to latent sources of SA against the unmodelled variance and identified a population of neurons whose shared SA variance exceeded the upper bound on the private variance (Figure 4k, neurons below the diagonal), indicating that the variability of these neurons arises primarily from common latent input.

### The statistics of evoked and spontaneous activity are consistent across individuals

We next sought to determine how representative our results were for a dataset of seven additional zebrafish larvae. Example fits for two of these fish are given in Supplementary Figures 6 and 7. For consistency of comparison between fish we again fit the statistical model with three latent factors, and characterised the distribution of correlation coefficients between the fluorescence trace and model fit by the mean and interquartile range (a description of variability that is more robust to outliers and skewness than, e.g. the standard deviation). Across the 8 fish the mean correlation coefficient was centered at ~ 0.6 (Figure 5a).

**Figure 5:**
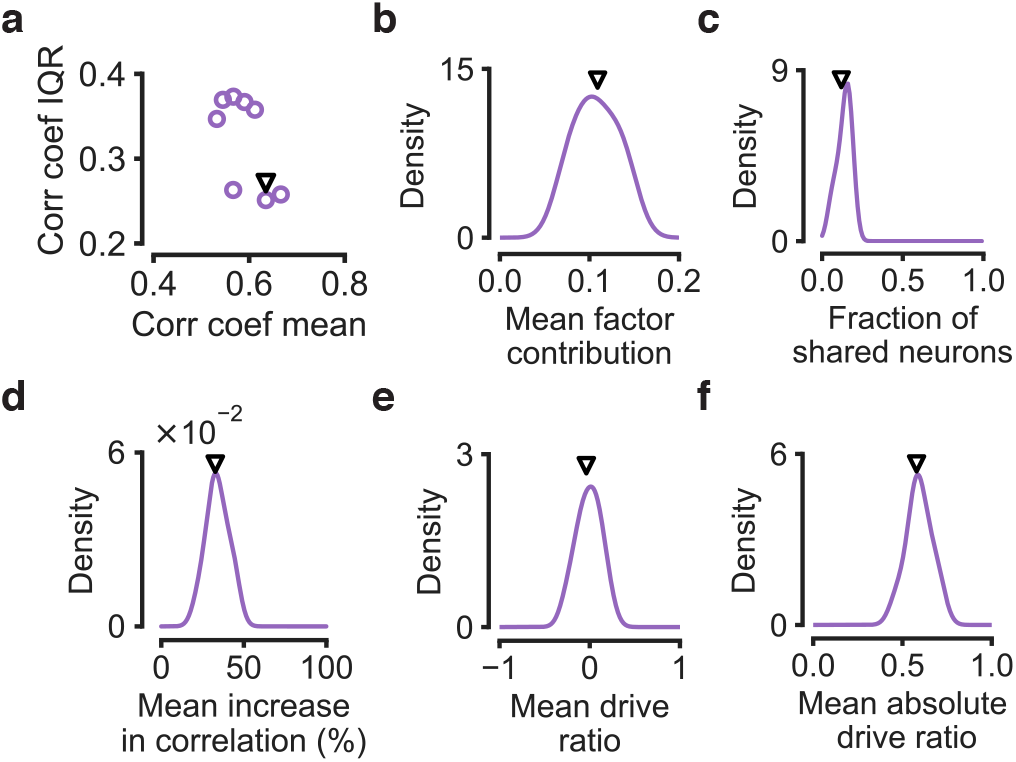
Summary of CILVA fits across a population of zebrafish larvae. Black triangles point to the example fish from Figure 1. **a**, Mean and interquartile range (IQR) of correlation coefficient distributions for *n* = 8 larvae. **b**, Distribution of factor contribution indices. For model fits with 3 latent sources of SA, each factor has a contribution index of ~ 0.1. **c**, Distribution of fraction of neurons ‘shared’ between multiple factors. Neurons were considered shared if they were coupled to more than one factor with coupling strengths exceeding a threshold of 25% of the maximum coupling strength for that factor. **d**, Mean improvement in correlation coefficients with incorporation of latent sources of SA (cf. Figure 4h,i). **e**, Distribution of mean drive ratios across the population of larvae. **f**, The mean absolute values of the drive ratio are greater than 0, showing that individual neurons tend to be biased towards either EA or SA. Histograms in panels **b** - **f** obtained by non-parametric density estimation with Gaussian kernels. Raw data points used for histograms given in Supplementary Table 3.

How variable is the influence of latent sources of SA across fish? We found that latent factors had average individual contribution indices of 0.1 (Figure 5b). Factors were also mostly nonoverlapping, with only a small fraction of neurons participating in multiple factors (Figure 5c). Incorporating all three factors increased the mean correlation coefficient between raw fluorescence data and model-fit by 34% on average compared to model-fits without SA (Figure 5d).

We then measured the relative influence of SA and EA in terms of the drive ratio. The distribution of mean drive ratios was centered at −0.01, suggesting that SA and EA were largely balanced within individual fish (Figure 5e). Our example fish was only marginally more in favour of SA than the other fish (mean drive ratio −0.04). Despite this balance, the set of drive ratios might all take values near zero (i.e., if every neuron in the population were driven equally by EA and SA), near one (the more extreme case, where every neuron is driven by either the stimuli or latent factors, but not both), or something in between. To discriminate between these possibilities we considered the absolute values of the drive ratios. The means were similar across fish (Figure 5f) with the distribution centered at 0.6. This indicates that, while EA and SA tended to be balanced at the population level, individual neurons mostly biased their activity towards being either stimulus-driven or spontaneous.

### Application to visual cortex

We next explored the application of the model to publicly available data from mouse primary visual cortex [27]. In this case, stimuli of higher dimension were presented more rapidly than in our previous application. Briefly, head-fixed mice expressing the calcium indicator GCaMP6s (via viral injection) stood on an air-suspended ball while drifting gratings were presented across the visual field with 1 to 3 second intervals and at 8 orientations, 3 spatial frequencies, and 4 temporal frequencies. This required a stimulus filter with 96 dimensions (compared to 9 in the zebrafish case). We verified with simulated data that the model could accurately recover evoked and spontaneous components in this regime (Supplementary Figure 2, Supplementary Table 2). We used a penalised regression approach to estimate the rise and decay time constants of the calcium transients given the stimulus presentation times (Methods), and then applied CILVA to decouple the evoked and spontaneous fluorescence components (Figure 6a). As this data consisted of raw fluorescence levels (rather than Δ*F*/*F*), the model parameters *β_n_* corrected for the baseline fluorescence. The model reconstruction provided a good fit, with correlation coefficients much larger than in the case of shuffled data (Figure 6b, Supplementary Figures 8–10). Extracted latent factors were more active than for the zebrafish larvae (Figure 6c), but were still largely non-overlapping (Figure 6d).

**Figure 6:**
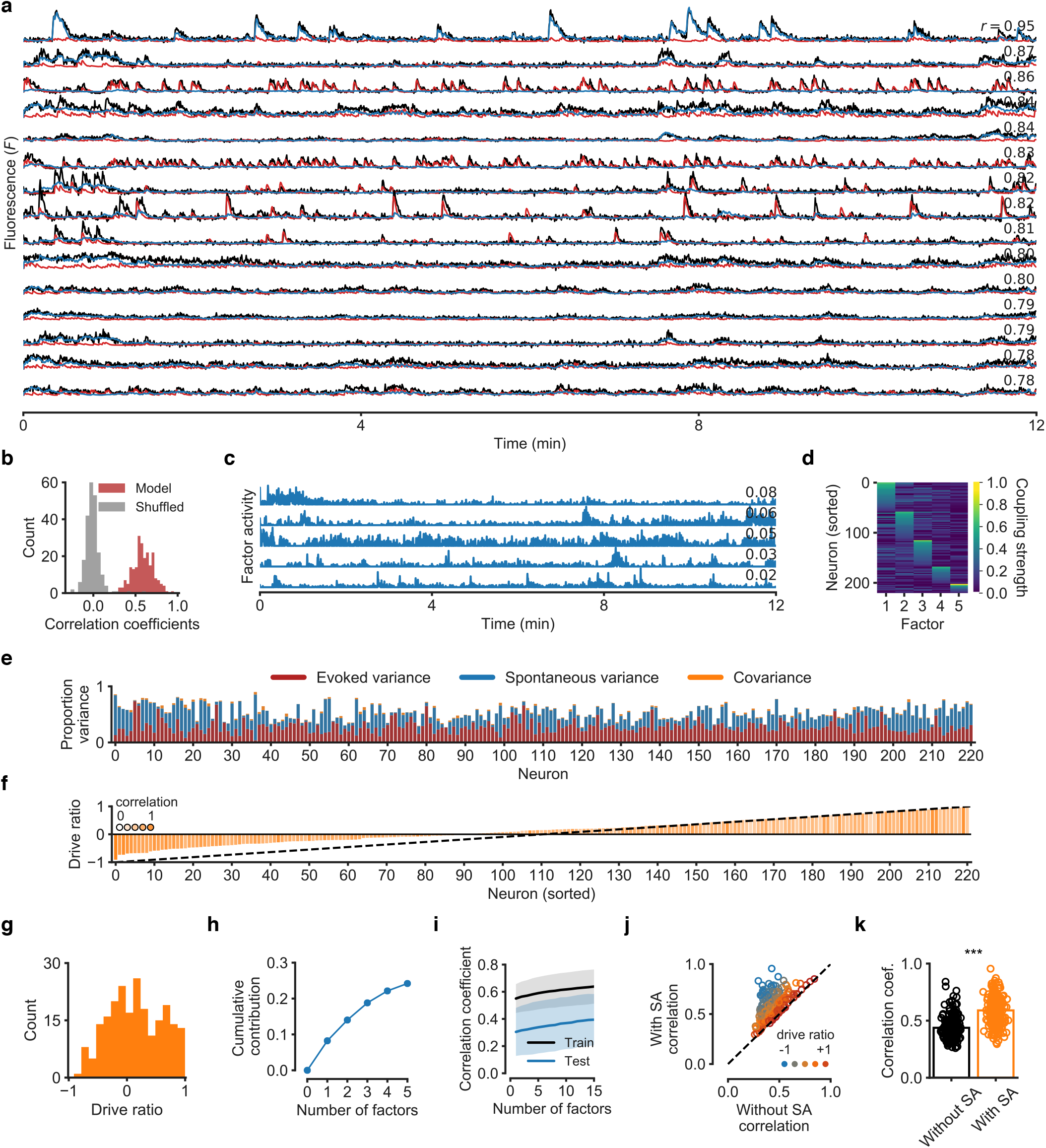
Application of CILVA to mouse visual cortex. **a**, Decoupled fluorescence traces showing evoked (red) and spontaneous (blue) components of neural activity from visual cortex. Inset numbers denote the Pearson correlation coefficient between raw fluorescence trace and model fit. **b**, Distribution of correlation coefficients between data and model fits. Shuffled data obtained by cyclically permuting each trace by a random offset while preserving its temporal structure. **c**, Inferred latent factor timeseries. Inset numbers denote the factor contribution index (Methods). **d**, Learned factor coupling matrix showing that the inferred factors target largely non-overlapping sets of neurons. **e**, Decomposition of each neuron’s sample variance in terms of variance attributable to EA, SA, and their interaction. In contrast to the zebrafish data, the mouse data suggests that individual neurons largely balance EA and SA. **f**, Drive ratios for a population of visual cortical neurons. **g**, The distribution of drive ratios exhibit a central mode at ~ 0, in contrast to the zebrafish data, indicating that for a model fit with 5 latent factors more neurons balance EA and SA. **h**, Cumulative contribution indices for 0 to 5 factors. **i**, Correlation coefficients show progressive improvement with increasing numbers of factors on both training data and held-out test data. **j**, Correlation coefficients between raw fluorescence trace and model fit with and without the SA component. Neurons with negative drive ratios (blue circles) demonstrate substantial improvement in the quality of model fit when incorporating SA. **k**, Improvement in the quality of model fit when incorporating the SA component is statistically significant (*p* < 0.001, Wilcoxon signed-rank test).

Decomposition of each neuron’s fluorescence trace in terms of the evoked and spontaneous components (Figure 6e) and drive ratio (Figure 6f), however, showed a qualitatively different effect than in the case of the zebrafish. There were more neurons with drive ratios close to zero compared to the zebrafish data (Figure 6g), indicating more balanced contributions from EA and SA at the single-neuron level. We fit the model with 5 latent factors. While the contribution indices for the factors gradually diminished (Figure 6h), varying the number of factors from 1 to 15 did not identify a point at which the overall quality of fit failed to increase, including in held-out test data (Figure 6i). However, similar to the zebrafish data, many neurons showed a substantial improvement in the quality of model fit when accounting for shared sources of SA (Figure 6j, neurons above the diagonal), with a statistically significant improvement of the average correlation coefficient across the population (Figure 6k).

## Discussion

Neural activity elicited in response to a stimulus can be substantially affected by ongoing SA. The CILVA approach for decoupling these influences has the advantage over simpler approaches, such as the sequential application of non-negative least squares and NMF, since receptive fields are inferred simultaneously with latent factors, preventing the latter from confounding measurements of the stimulus-evoked response. Not only does this allow us to estimate tuning curves that are unbiased by spontaneous calcium transients, but also to estimate the latent structure of SA alone, unbiased by evoked responses. The composition of a neuron’s sample variance can then be straightforwardly expressed in the model in terms of the variance of the decoupled evoked and spontaneous components, together with their covariance. CILVA thus provides a new tool for quantitative analyses of the interaction between EA and SA in single trials, reducing dependence on approaches to sensory coding that require averaging away potentially important information encoded in SA.

Many analyses of calcium imaging data deconvolve calcium transients to estimate the underlying neural activity before using more traditional methods of spike train analysis. Here we jointly modelled the underlying sources of activity together with the calcium transients themselves, allowing a direct comparison between the raw imaging data and the model components, and avoiding the intermediate computational step of deconvolution, which can impact model performance compared to joint inference approaches (see e.g. [14]). However, CILVA is closely related to latent factor models for spike train data. Gaussian-process factor analysis, for example, assumes that population spiking activity is linearly driven by a small number of latent factors evolving smoothly through time according to a Gaussian process [9]. A similar model, the Poisson linear dynamical system, models neural activity by Poisson processes, where firing rates across the population are driven by a hidden low dimensional linear dynamical system [10]. These models consider a neuron to be a noisy sensor of an underlying latent state, and the smooth path that population activity traces through this low dimensional state space constitutes the underlying computation implemented by a neural circuit [28]. In contrast to such models, which explicitly constrain the temporal evolution of latent factors, our statistical model assumes that latent factor activity states at each time point are independent and identically distributed according to a (non-negative) maximum entropy prior. Autocorrelation of the latent factors then arises due to their convolution with a calcium impulse response function. While an explicit dynamics could be imposed on the latent factors [14], we chose not to do so due to a conflict of timescales: the relevant neural dynamics often takes place over several hundred milliseconds [9], but this may only constitute a few imaging frames in calcium imaging data. Thus, calcium transients predicted by the model may appear erroneously prolonged if factor activity states could only change gradually.

Additive interactions between EA and SA, as assumed by the model, have been identified in numerous studies. For example, optical imaging of cat visual cortical neurons using voltage-sensitive dyes [1], and cellular-resolution two-photon calcium imaging [2], multiple simultaneous Neuropixels probes [2], and wide-field calcium imaging of both cortical hemispheres [29] in mouse visual cortex have all shown substantial additive modulation of evoked responses by coordinated SA that proceeds unimpeded by stimulus onset. However, there may be cases where the interaction between EA and SA is more complex than a simple additive scheme. For example, trial-to-trial variability of evoked responses could result from changes in excitability, reflecting a multiplicative effect of SA. Such an interaction could potentially be included in our model by incorporating an appropriate nonlinear activation function (similar to ref. [30]).

We identified subsets of neurons whose SA primarily originated from shared sources, with the SA of the remaining neurons arising either from private sources or from residual imaging noise. In contrast with auditory cortex, where it has been suggested that fluctuations of subthreshold membrane potentials arise primarily by shared sources [31], our analysis suggests that in the optic tectum a significant fraction appear to be driven primarily by private, as opposed to shared, SA. Nevertheless, shared SA was responsible for a substantial portion of the variance across the population, and the model accounting for shared SA significantly outperformed the model that did not. Moreover, this fraction represents merely a lower bound on the number of neurons dominated by shared SA, and this bound could increase as the recorded set of tectal neurons approaches the complete population. Indeed, SA that we currently consider private may in reality be shared, but due to experimental constraints in the optical imaging system we may simply have not observed the neurons with similar profiles of SA. Okun et al. [32] found that the correlation structure of cortical populations in primates and mice could be well-predicted by the coupling of individual neurons to the population firing rate, a one dimensional measure of activity. ‘Choristers’ have firing rates coupled to the population, and are thus dominated by shared variability, whereas ‘soloists’ are less affected by population-wide events and are dominated by private variability, even during SA. In our zebrafish data, in contrast, population activity was best described by coupling of neurons to one of multiple latent factors, and these factors could not be described by a single latent state governing SA since they were mutually independent.

Our method identified multiple independent latent sources of SA targeting distinct, largely nonoverlapping sets of neurons. Even a primary sensory area like the optic tectum or visual cortex receives converging inputs from other brain regions that can make it highly active in the absence of sensory inputs [26, 33, 34]. Recently, several studies reported brain-wide activity correlated with behaviour [2, 5, 35]. For example, Stringer et al. analysed calcium imaging data from 10, 000 neurons in the mouse primary visual cortex and found that locomotor variables such as pupil diameter and running speed accounted for ~ 20% of the total variance of the population activity [2]. Potentially, similar inputs to the tectum for the purpose of, e.g., visuomotor integration [17], could form the physiological basis of the latent factors that we extracted. However, while overt behavioural parameters like pupil diameter can be unambiguously measured and correlated with neural activity, CILVA attempts to adapt to any kind of input that induces structured patterns of SA that linearly combine with stimulus-evoked responses, even if such inputs are not directly measured.

## Methods

### Zebrafish

All procedures were performed with approval from The University of Queensland Animal Ethics Committee (approval certificate number QBI/152/16/ARC). *Nacre* zebrafish (*Danio rerio*) embryos expressing *elavl3*:H2B-GCaMP6s, of either sex, were collected and raised according to established procedures [36] and kept under a 14/10 hr on/off light cycle.

Zebrafish larvae were embedded in 2.5% low-melting point agarose, positioned at the centre of a 35 mm diameter plastic petri dish and overlaid with E3 embryo medium. Calcium imaging was performed at a depth of 70 *μ*m from the dorsal surface of the tectal midline. Time-lapse two-photon images were acquired using a Zeiss LSM 710 inverted two-photon microscope. A custom-made inverter tube composed of a pair of beam-steering mirrors and two identical 60 mm focal length lenses arranged in a 4f configuration was used to allow imaging with a 40X/1.0 NA water-dipping objective (Zeiss) in an upright configuration. Samples were excited via a Spectra-Physics Mai TaiDeepSee Ti:Sapphire laser (Spectra-Physics) at an excitation wavelength of 940 nm and the emitted light was bandpass filtered (500–550 nm). Laser power at the sample ranged between 12 to 20 mW. Images of 416×300 pixels were obtained at 2.1646 Hz. To improve the stability of the recording, chambers were allowed to settle for three hours prior to start of two-photon imaging.

Visual stimuli were projected on white paper placed around the wall of a 35 mm diameter petri dish using a projector (PK320 Optoma, USA), covering a horizontal field of view of 174°. A red filter (Zeiss LP590 filter) was placed in the front of the projector to avoid interference of the projected image in the signal collected by the detector. Larvae were aligned with one eye facing the white paper side of the dish and with the body axis orthogonal to the projector. Visual stimuli were generated using custom software based on MATLAB (MathWorks) and Psychophysics Toolbox. Each trial consisted of 6° diameter black spots at nine different positions, separated by 15° intervals from 45° to 165°, where 0° was defined as the direction of the larva’s body axis. Their order was set to maximise spatial separation within a trial (45°, 120°, 60°, 135°, 75°, 150°, 90°, 165°, 105°). Spots were presented for 1 s, followed by 19 s of blank screen. We projected consecutive trials of nine spots with 25 s of inter-trial interval.

### Mice

We used data available at ref. [27]. The experimental procedures are described in detail in refs. [37, 38]. Neurons were recorded simultaneously at 2.5Hz using calcium imaging. Visual stimuli were shown at approximately 1 Hz, with randomized inter-stimulus intervals. The stimuli were drifting gratings with 8 directions, 4 spatial frequencies and 3 temporal frequencies. Blank stimuli (gray screen) were also interleaved. While this data was originally recorded from multiple imaging planes, to avoid issues with timing of latent factor activity we selected neurons from a single imaging plane. We then max-normalised the amplitude of the fluorescence traces, and subsampled neurons with variance exceeding 1.5 × 10^−3^, reducing the population size to *N* = 221 of the most active neurons.

### Residual NMF method

We defined stimulus regressors *ϕ_i_* ∈ ℝ^*T*^ for *i* = 1,…, *K* by convolving the binary stimulus time-series **s**_*i*_ ∈ ℝ^*T*^ with a calcium impulse response kernel **k** (a difference-of-exponentials function, defined below), giving *ϕ_i_* = **k** * **s**_*i*_. We then estimated regression coefficients *β_ni_* by solving the non-negative minimisation problem

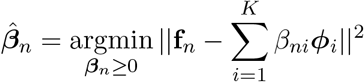

where ***β***_*n*_ = (*β*_*n*1_,…, *β_nK_*)^⊤^. The evoked component of the fluorescence signal can be defined in terms of the estimated regression coefficients as

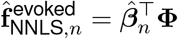

where 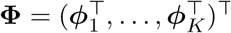. Here NNLS refers to the non-negative least squares algorithm used to perform the minimisation. Residual data **e**_*n*_ was then defined as

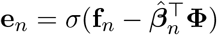

where *σ* is a linear rectifier *σ*(*x*) = max(0, *x*) applied elementwise ensuring non-negativity of the residuals. We then applied NMF to the residual data 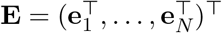 by solving the minimisation problem

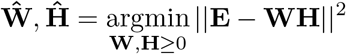

where **W** ∈ ℝ^*N,L*^ and **H** ∈ ℝ^*L,T*^. The *L* rows of **Ĥ** are timeseries describing the evolution of low dimensional structure, and the columns of **Ŵ** describe how neurons are coupled to such timeseries. The NMF-estimated SA for neuron *n* is given by the projection of the latent timeseries onto a single dimension by the corresponding row of **Ŵ**

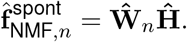

The full fluorescence data can thus be approximately reconstructed as

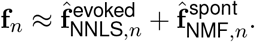

We use the NNLS routine in SciPy and the NMF routine in scikit-learn [39].

To evaluate the models of SA produced by NMF we then expressed the spontaneous components in terms of a basis of calcium impulses by solving the non-negative minimisation problem

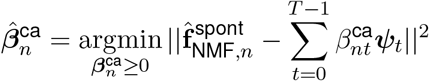

where each ***ψ***_*t*_ is a calcium response following a unit impulse 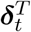 at time *t*,

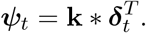

Here 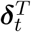 is a vector of length *T* that takes the value of 1 at *t* and 0 elsewhere. The expression of 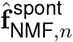 in the basis of calcium transients is then given by 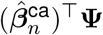, where 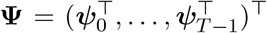. Note that this differs from the basis of stimulus regressors used to model the stimulus-driven component of neural activity as we employ a regressor for each time point *t* in the entire trace.

### CILVA model

#### Fluorescence model

We model fluorescence data *f_n_*(*t*) as a linear transformation of the calcium concentration *c_n_*(*t*) plus independent and identically distributed additive Gaussian noise. The generative model for the observed fluorescence of neuron *n* is thus

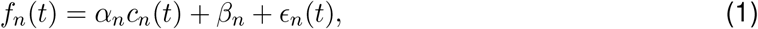

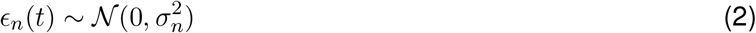

where *α_n_* is a scaling factor and *β_n_* is the baseline fluorescence level of neuron *n*. The assumption of Gaussian noise is a simple and tractable way to account for noise in both the calcium concentration and noise due to optical imaging. This model is standard for fluorescence imaging data [7, 20, 40].

#### Calcium dynamics

The calcium concentration *c_n_*(*t*) is generated as the convolution of a difference-of-exponentials kernel **k** with a function ***λ***_*n*_ that determines the intensity of neural activity,

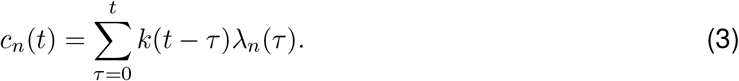

The kernel **k** captures the stereotypical rise-and-decay calcium dynamics, which are assumed to possess time constants that are unchanging throughout the recording

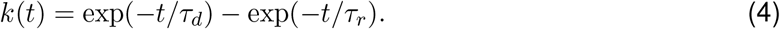

An explicit rise time was essential for modelling the experimental data with a GCaMP6s calcium indicator [6]. For the zebrafish data used in the paper we used calcium transient time constants of *τ_r_* = 5.68/*F_s_* and *τ_d_* = 11.5/*F_s_*, where *F_s_* = 2.1646 Hz is the imaging rate of the fluorescence microscope. For the mouse recording we estimated the time constants using a penalised regression approach described in the Model Fitting section below.

#### Intensity function

Changes in the intracellular calcium concentrations are driven by an intensity function ***λ***_*n*_ for each neuron *n*. We take advantage of the fact that we expect evoked responses to be time-locked to the presentation of a stimulus, with the remaining signal attributable to structured SA. The intensity is thus comprised of a stimulus drive and a latent drive

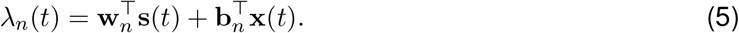

Here **w**_*n*_ ∈ ℝ^*K*^ corresponds to the stimulus filter for neuron *n*, **s**(*t*) ∈ ℝ^*K*^ is the vector describing which stimulus is active at time *t* (under a 1-of-*K* encoding scheme, such that **s**(*t*) has a 1 in index *i* if the *i*th stimulus is active, and zeros elsewhere), **b**_*n*_ ∈ ℝ^*L*^ is a row of the factor loading matrix, and **x**(*t*) ∈ ℝ^*L*^ is the activity level of the latent factors at time *t*. Thus, at each time point the latent drive is the projection of a low dimensional latent process **x**(*t*) into one dimension. By fixing the onset of the stimulus drive and leaving the latent factors unconstrained, we allow the factors to adapt to the patterns of SA in the data.

#### Latent factors

The factors **x**(*t*) exist in a latent space that is low dimensional relative to the population size *N*. Consequently, the variability that they account for in the model must be shared among groups of neurons. We place an exponential prior on the latent factors to encourage sparsity,

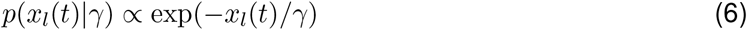

for 0 ≤ *t* ≤ *T* − 1 and 1 ≤ *l* ≤ *L*. Here *γ* is the parameterisation of the exponential distribution in terms of its mean; i.e., 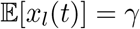, which acts as a sparsity penalty. Selection of *γ* is described in the Model Fitting section below.

#### Evoked and spontaneous variance components

Given the fitted model parameters, we defined the evoked and spontaneous components of the fluorescence signal as

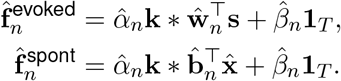

Note that we include the baseline fluorescence term 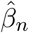 to ensure the evoked and spontaneous traces are appropriately aligned with the raw fluorescence signal during visual comparisons. The variance of the reconstructed fluorescence levels can then be written in terms of these components

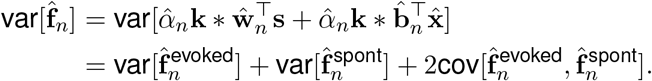

When plotting variance components as proportions of sample variance as in Figure 4a, the sample variance var[**f**_*n*_] is corrected for imaging noise by subtracting the estimated sample imaging noise variance 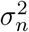. We then define the drive ratio for neuron *n* using the variance components as

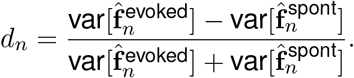

This defines an index ranging from −1 to 1 that describes the extent to which a neuron is driven more by shared sources of SA or by EA.

#### Factor contribution index

The contribution of a factor **x**_*l*_ is defined as the average reduction in explained correlation caused by removing factor *l* from the model reconstruction of the fluorescence trace,

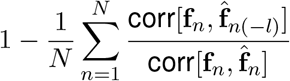

where 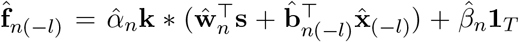, and 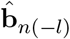 and 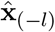 are obtained by deleting element *l* and row *l* from 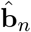 and 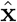, respectively.

#### Tuning curve comparison

Tuning curves obtained by averaging were defined as the mean Δ*F*/*F* over the 4th to 7th frames following stimulus onset. As the stimulus filters {**ŵ**_*n*_} are rescaled by our parameter identification algorithm (described below), we compared the averaging-based tuning curves with 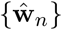, where *k*_max_ = max_*t*_ *k*(*t*) is the maximum value of the calcium kernel. This scaling of the stimulus filter reports the amplitudes of the calcium transients evoked by each stimulus, which are directly comparable with the tuning curves obtained by averaging.

### Model fitting

#### Maximum a posteriori inference

We fit the model by maximising the posterior density of the latent variables,

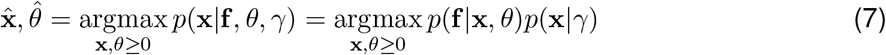

where the parameters of the model are *θ* = ({*α_n_*}, {*β_n_*}, {**w**_*n*_}, {**b**_*n*_}). Ideally, one could perform this optimisation using the expectation-maximisation algorithm, which alternates between computing the posterior distribution over the latent factors *q*(**x**) = *p*(**x**|**f**, *θ, γ*), and maximising the posterior expectation 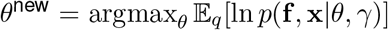. However, the E-step is not analytically tractable since our exponential prior on *x_l_*(*t*) is non-conjugate for the likelihood model. Instead, we use a related approach that alternately optimises Equation 7 according to the steps

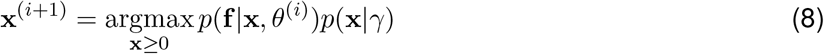

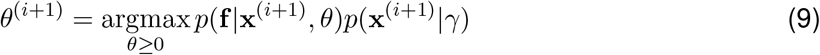

until numerical convergence or until *i* reaches a user-specified number of iterations. The alternating maximisations are each performed using the bounded BFGS algorithm with limited memory (L-BFGS-B), with exact gradients derived below.

The logarithm of the joint model probability density is

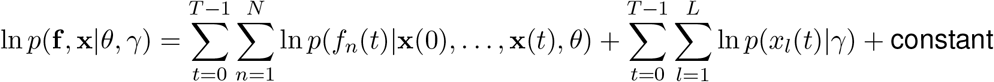

where the constant term does not depend on the parameters of *θ* to be estimated. Let *ℓ*(**x**, *θ*) = ln *p*(**f, x**|*θ, γ*), and let 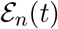 denote the model reconstruction error for neuron *n* in imaging frame *t*,

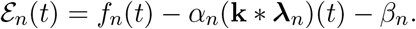

The derivatives of *ℓ*(**x**, *θ*) with respect to the parameters and latent variables are then

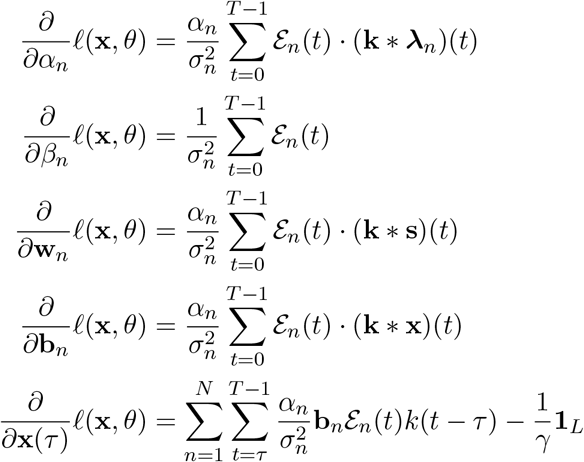

where for a matrix **Λ** ∈ ℝ^*T*×*q*^ the convolution **k** * **Λ** ∈ ℝ^*T*×*q*^ is performed row-wise. In practice we vectorise the computation of the gradients to improve efficiency.

The imaging noise variance terms 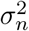 are estimated using the method in ref. [20]. Specifically, 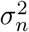 is estimated as the mean of the power spectral density of **f**_*n*_ over the range (*F_s_*/4, *F_s_*/2), where *F_s_* is the imaging rate of the fluorescence microscope.

#### Model identifiability

As is common in factor analysis-style methods, the model parameters and latent variables (*θ*, **x**) are not uniquely identifiable in their current form. Our model-fitting algorithm thus transforms the estimates 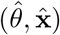 into a standardised form according to the following procedure. First we fit the CILVA model to data {**f**_*n*_} using the MAP estimator to obtain model parameters 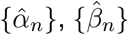, {**ŵ**_*n*_}, 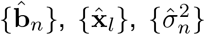. We then sort factors 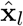 in descending order of their Euclidean norm so that 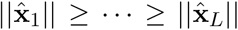, and sort factor coupling column vectors 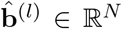 to take the same order. Next, we normalise latent factors and proportionally rescale factor coupling vectors,

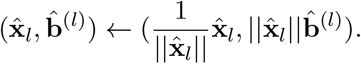

The latent factors are now identifiable. Finally, we normalise the static model parameters by the norm of the neural intensity vector,

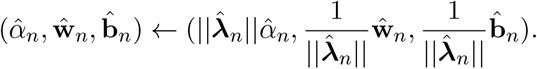

This ensures identifiability of the static model parameters *θ*.

#### Parameter initialisation

For the application of CILVA to mouse data, we implemented a simple penalised regression approach to estimate the calcium transient time constants *τ_r_* and *τ_d_*. The idea is to alternately estimate tuning curves (using knowledge of the stimulus presentation times) and update our time constants given these new tuning curves. The constants *τ_r_* and *τ_d_* must respect the inequality

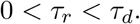

We thus parameterise *τ_d_* in terms of the rise time constant and a positive offset,

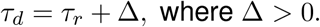

Let *k*_*τ_r_*,Δ_(*t*) = exp(−*t*/(*τ_r_* + Δ)) − exp(−*t*/*τ_r_*). Given some values of *τ_r_* and Δ, we define **Φ**_*τ_r_*,Δ_ ∈ ℝ^*K*×*T*^ analogous to the residual NMF method with

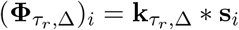

for *i* = 1,…, *K*. For every neuron *n* we then fit tuning curves as

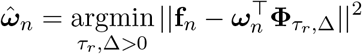

using non-negative least squares. Then, given a set of tuning curves 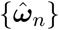, we update the time constants by minimising the model reconstruction error averaging over all neurons,

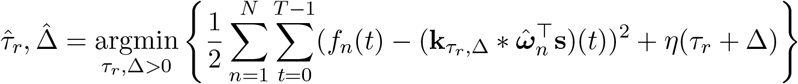

where *η* > 0 is a chosen penalty coefficient. The derivative of this term with respect to 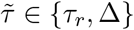 is

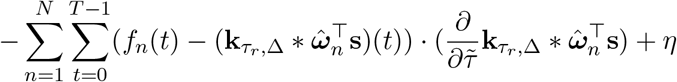

where

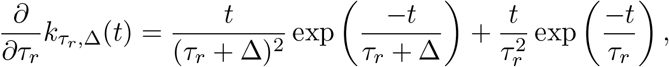

and

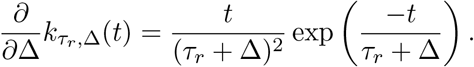

We perform the non-negative minimisation with these gradients using L-BFGS-B. Learning the time constants typically only required several alternations of estimating the tuning curves 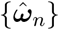 and updating the time constants 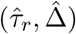.

We use the stimulus regressors to also initialise the filters **w**_*n*_; i.e.,

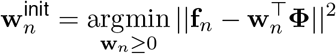

with the calcium time constants obtained either by the penalised regression approach described above or by manual specification. We then initialise *α_n_* as a small perturbation around 1, 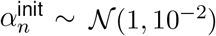, and 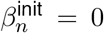. We initialise the latent factor coupling strengths as uniform samples from the unit interval, 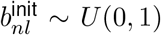, and the factor activity levels uniformly from a small interval, 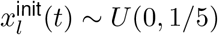.

#### Model selection

CILVA depends on two key hyperparameters: the number of latent factors *L* and the sparsity parameter *γ*. Here we describe how we estimate these hyperparameters. To avoid local minima, we fit the model several times to the data with different random initialisations of the factor coupling vectors {**b**_*n*_} and factor activities {**x**_*l*_}. For a given *L* and *γ* we fit the latent variables and parameters on training data as

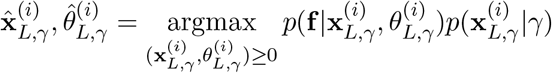

where *i* = 1,…, *i*_max_ denotes the *i*th initialisation of **x** and *θ*. We select the optimal parameters 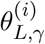 and hyperparameter *γ* as those that maximise the joint density of the data and latent variables on 5 minutes of held-out test data **f**^test^; i.e.,

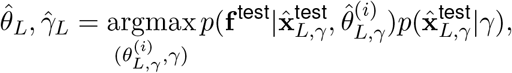

where values of *γ* are obtained via grid search over a small interval [Δ_*γ*_, *d*Δ_*γ*_] with step-size Δ_*γ*_ and number of grid points *d*. Here we infer new latent variables 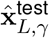 that explain the patterns of spontaneous activity in the test data **f**^test^. The selected latent variables for the training data are then those that correspond to the optimal *θ* and *γ*,

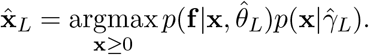

We used Δ_*γ*_ = 0.2, *d* = 10 and *i*_max_ = 5. For the example zebrafish used in the main text we selected *L* as the number of factors after which the mean correlation coefficient between the raw fluorescence traces and model-reconstructions failed to substantially improve (i.e., at the ‘elbow’ in Figure 3d).

Fitting CILVA for testing and preliminary data analysis required a computation time of ~10 minutes on a 64-bit MacBook Pro with a 3.1 GHz Intel Core i7 Processor and 8 GB DDR3 RAM running Python 3.6.4. For the model fits in this paper we allowed the optimisation procedure to run to a user-specified number of alternations of Equation 8 and Equation 9 (typically ~100), performed on a computer cluster with 17 Dell EMC PowerEdgeR740 compute nodes, each comprised of two Intel Xeon Gold 6132 processors with 384 GB DDR4 RAM. Scheduled jobs were allocated 2 CPUs and 5GB RAM, and required ~1 hour to complete.

#### Simulated data

To generate simulated data we sampled the latent factors from a zero-inflated exponential distribution with probability *ξ* of a non-zero latent event,

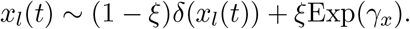

This ensured the latent factor activity was sparse. We also introduced a private SA term *z_n_*(*t*) for neuron *n* at time *t* by sampling from a zero-inflated exponential with probability *π* of a non-zero private event,

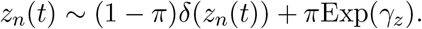

The intensity function was then given by

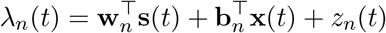

with the fluorescence levels following the standard CILVA model with a common imaging noise variance *σ*^2^,

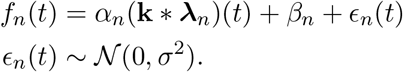

We sampled *α_n_* from the discrete uniform distribution on {2,…, 10} and for simplicity set *β_n_* = 0. The tuning curves **w**_*n*_ were defined as Gaussian functions *x* ↦ exp(−(*x* − *μ_n_*)^2^/2*ν*). For the simulation of data in response to well-spaced, low dimensional stimuli (cf. Figure 3) we sampled the centres *μ_n_* uniformly from the interval [0, *K*], where *K* is the number of stimuli, and sampled the widths *ν_n_* uniformly from [0, *K*/2]. For the simulation of data in response to rapidly presented, high dimensional stimuli (cf. Figure 6) we chose our receptive fields to be more selective and sampled *ν_n_* uniformly from 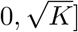.

Factor coupling vectors **b**_*n*_ were defined by evenly assigning the *N* neurons to *L* factors, and sampling *b_nl_* ~ *U*[*q*, 1] if neuron *n* is assigned to factor *l*, and *b_nl_* ~ *U*[0, 1 − *q*] otherwise. We found *q* = 0.85 provided simulations that appeared similar to the experimental data. We characterised the model reconstruction quality in terms of *π* and *σ*^2^ in Supplementary Figures 1 and 2, with the associated model parameters provided in Supplementary Tables 1 and 2.

## Acknowledgements

We thank Carsen Stringer for helpful feedback on an earlier version of the paper. This work was supported by Australian Research Council Discovery Projects 170102263 and 180100636 awarded to G.J.G. M.A.T. is supported by an Australian Government Research Training Program Scholarship. Imaging was performed at the Queensland Brain Institute’s Advanced Microscopy Facility using an LSM 710, supported by the Australian Government through the ARC LIEF grant LE130100078.

## Data availability

Code for fitting the CILVA model and data for the example zebrafish in Figures 1–4 are available at https://github.com/GoodhillLab/CILVA. Data used for Figure 6 are available at ref. [27].

## Supplementary material

**Supplementary Table 1:** Parameters for simulated data corresponding to the presentation of a low dimensional stimulus with prolonged interstimulus intervals (analogous to the zebrafish data). Parameters related to time are defined with respect to imaging rate. Listed values of *π* and *σ*^2^ are defaults, but are varied over the range specified in parentheses.

| Parameter | Interpretation | Value |
| --- | --- | --- |
| $N$ | Number of neurons | 200 |
| $T$ | Simulation duration (frames) | 2000 |
| – | Imaging rate (Hz) | 2.1646 |
| $\tau_r$ | Calcium rise time constant (frames) | 2.6 |
| $\tau_d$ | Calcium decay time constant (frames) | 5.3 |
| $K$ | Number of stimuli | 10 |
| – | Stimulus repetitions | 5 |
| – | Interstimulus interval (frames) | 39 |
| $L$ | Number of latent factors | 4 |
| $\xi$ | Probability of latent factor event (per frame) | 0.1 |
| $\pi$ | Probability of private spontaneous event (per frame) | 0.05 (0.0 - 0.3) |
| $\gamma_x$ | Mean intensity of latent event | 0.1 |
| $\gamma_z$ | Mean intensity of private spontaneous event | 0.1 |
| $\sigma^2$ | Imaging noise variance | 0.16 (0.04 - 0.4) |

**Supplementary Table 2:** Parameters for simulated data with rapid presentation of a high dimensional stimulus (analogous to the mouse data). Parameters related to time are defined with respect to imaging rate. Listed values of *π* and *σ*^2^ are defaults, but are varied over the range specified in parentheses.

| Parameter | Interpretation | Value |
| --- | --- | --- |
| $N$ | Number of neurons | 200 |
| $T$ | Simulation duration (frames) | 1800 |
| — | Imaging rate (Hz) | 2.5 |
| $\tau_r$ | Calcium rise time constant (frames) | 2.6 |
| $\tau_d$ | Calcium decay time constant (frames) | 5.3 |
| $K$ | Number of stimuli | 100 |
| — | Stimulus repetitions | 3 |
| — | Interstimulus interval (frames) | 6 |
| $L$ | Number of latent factors | 5 |
| $\xi$ | Probability of latent factor event (per frame) | 0.1 |
| $\pi$ | Probability of private spontaneous event (per frame) | 0.05 (0.0 - 0.3) |
| $\gamma_x$ | Mean intensity of latent event | 0.1 |
| $\gamma_z$ | Mean intensity of private spontaneous event | 0.1 |
| $\sigma^2$ | Imaging noise variance | 0.16 (0.04 - 0.4) |

**Supplementary Table 3:** Raw data points for the histograms in Figure 5. Zebrafish 5 corresponds to the example used in Figures 1–4 and Supplementary Figures 3–5.

|  | Zebrafish |  |  |  |  |  |  |  |
| --- | --- | --- | --- | --- | --- | --- | --- | --- |
|  | 1 | 2 | 3 | 4 | 5 | 6 | 7 | 8 |
| Correlation coefficient mean | 0.67 | 0.59 | 0.54 | 0.53 | 0.63 | 0.57 | 0.61 | 0.56 |
| Correlation coefficient IQR | 0.26 | 0.37 | 0.37 | 0.35 | 0.25 | 0.26 | 0.36 | 0.37 |
| Mean factor contribution | 0.14 | 0.10 | 0.10 | 0.08 | 0.11 | 0.12 | 0.13 | 0.07 |
| Fraction of shared neurons | 0.09 | 0.06 | 0.19 | 0.15 | 0.12 | 0.15 | 0.17 | 0.15 |
| Mean increase in correlation (%) | 44.02 | 30.16 | 30.17 | 34.76 | 32.70 | 35.88 | 41.56 | 23.41 |
| Mean drive ratio | -0.12 | 0.08 | 0.09 | 0.03 | -0.04 | -0.09 | -0.23 | 0.19 |
| Mean absolute drive ratio | 0.57 | 0.66 | 0.56 | 0.70 | 0.58 | 0.60 | 0.61 | 0.47 |

**Supplementary Figure 1:**
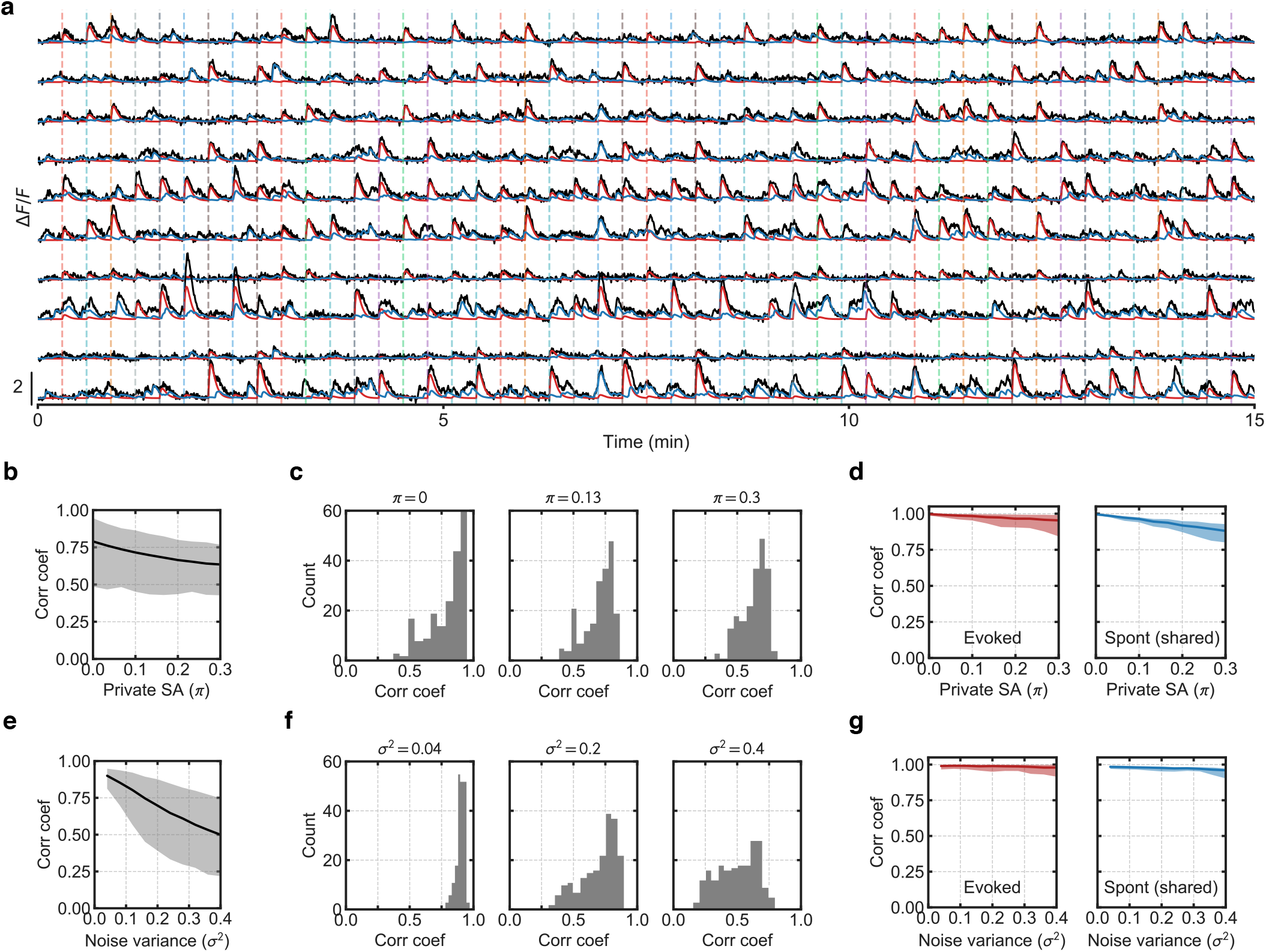
Results on simulated data (case 1). To validate performance we fit the model to simulated data (Methods). The two primary constraints on model performance are (i) the rate *π* of private spontaneous events, and (ii) the variance *σ*^2^ of the imaging noise. We thus systematically varied these two parameters and observed the ability of the model to recover the underlying evoked and spontaneous components. Parameters used in the simulations are given in Supplementary Table 1. **a**, Ten randomly chosen neurons from an example simulation with *π* = 0.05 and *σ*^2^ = 0.1. Black traces show simulated raw fluorescence data. The true composition of the fluorescence trace is given in red (EA) and blue (shared SA). **b**, The correlation coefficient between the raw fluorescence trace and model reconstruction decreases as the rate of private spontaneous events increases. **c**, Histograms of correlation coefficients for three example values of *π*. **d**, While the correlation coefficient decreases with *π*, recovery of the evoked (left) and spontaneous (right) fluorescence components remains highly accurate. **e-g**, Same as **b-d** but with varying noise variances *σ*^2^. High noise variances limit the correlation between the raw (noisy) fluorescence trace and (noiseless) model reconstruction, but recovery of the evoked and spontaneous components is still very robust. All shaded regions represent 95th percentiles.

**Supplementary Figure 2:**
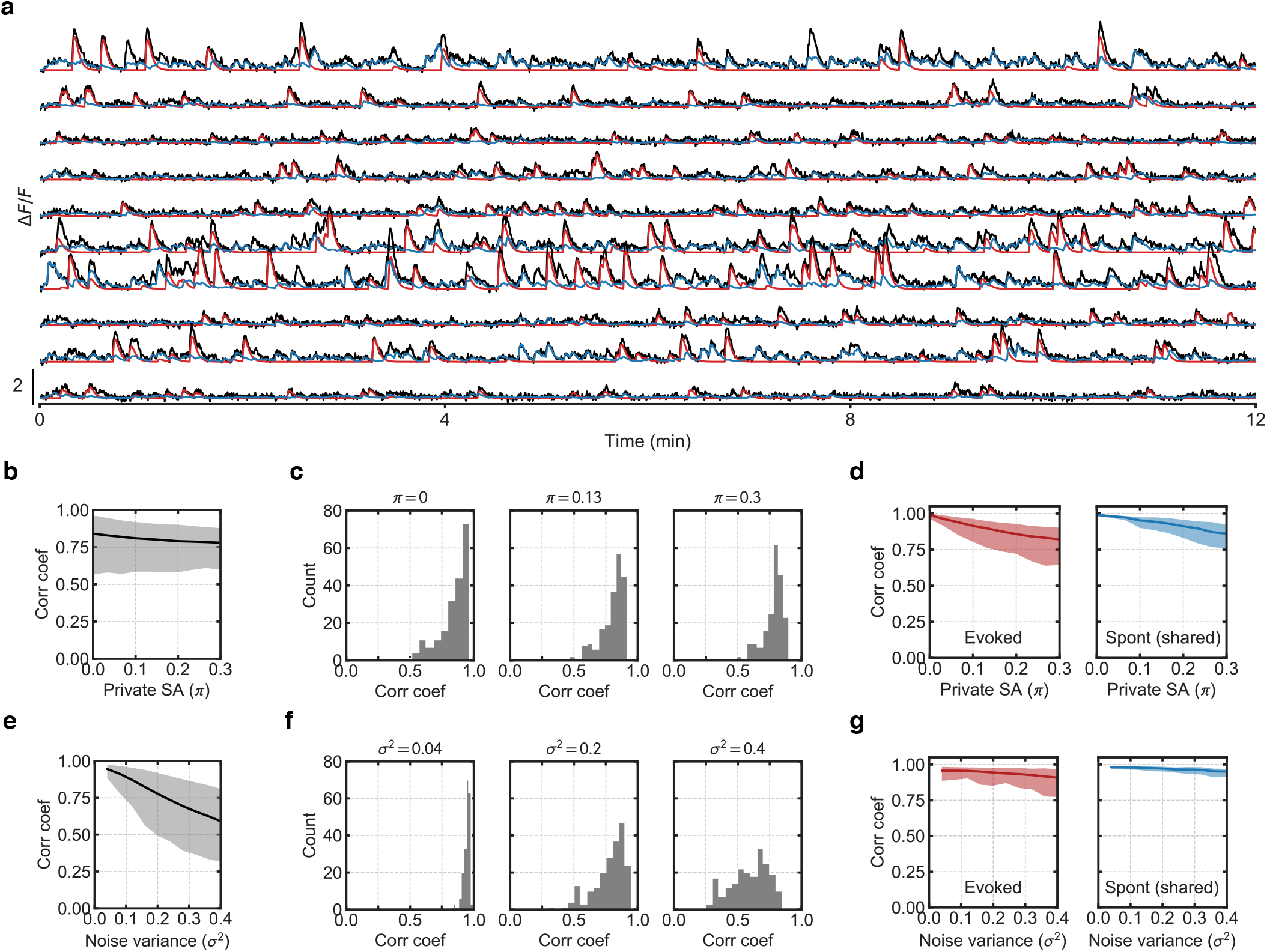
Results on simulated data with rapidly presented high-dimensional stimuli (case 2). Parameters used in the simulations are given in Supplementary Table 2. **a**, Ten randomly chosen neurons from an example simulation with *π* = 0.05 and *σ*^2^ = 0.1. Black traces show simulated raw fluorescence data. The true composition of the fluorescence trace is given in red (EA) and blue (shared SA). **b**, The correlation coefficient between the raw fluorescence trace and model reconstruction decreases as the rate of private spontaneous events increases. **c**, Histograms of correlation coefficients for three example values of *π*. **d**, While the correlation coefficient decreases with *π*, recovery of the evoked (left) and spontaneous (right) fluorescence components remains highly accurate. **e-g**, Same as **b-d** but with varying noise variances *σ*^2^. All shaded regions represent 95th percentiles.

**Supplementary Figure 3:**
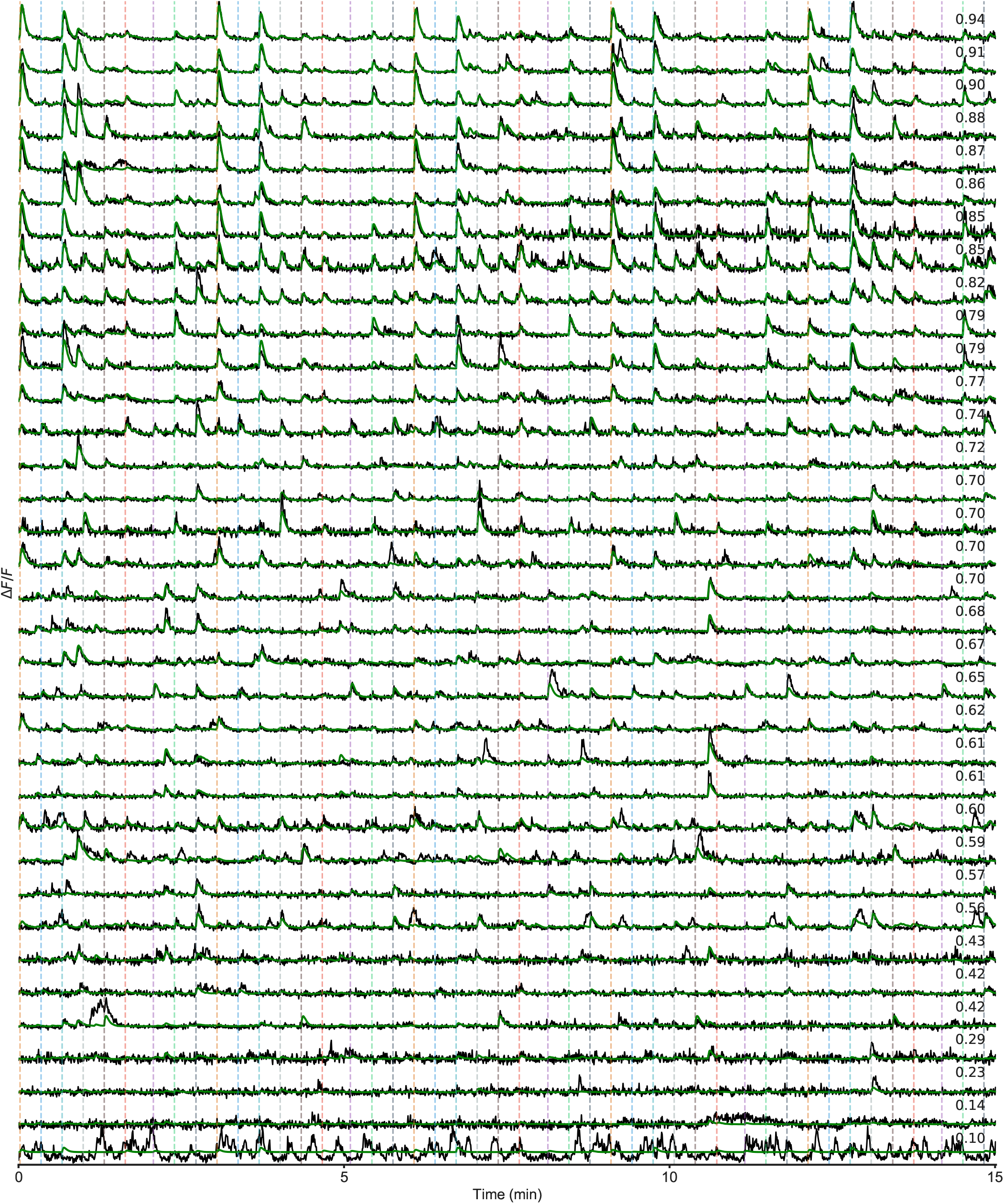
Model fits for 35 neurons uniformly sampled from the larval zebrafish in Figure 1. Example fluorescence traces (black) and corresponding model fits (green). Dashed vertical lines indicate stimulus onset times. Inset numbers denote Pearson correlation coefficient between raw trace and model fit. Sampled neurons are sorted by correlation. Poor fits can result from neurons that show inconsistent responses (or no responses) to presented stimuli or neurons dominated by private SA (and therefore that cannot be assigned to a latent factor).

**Supplementary Figure 4:**
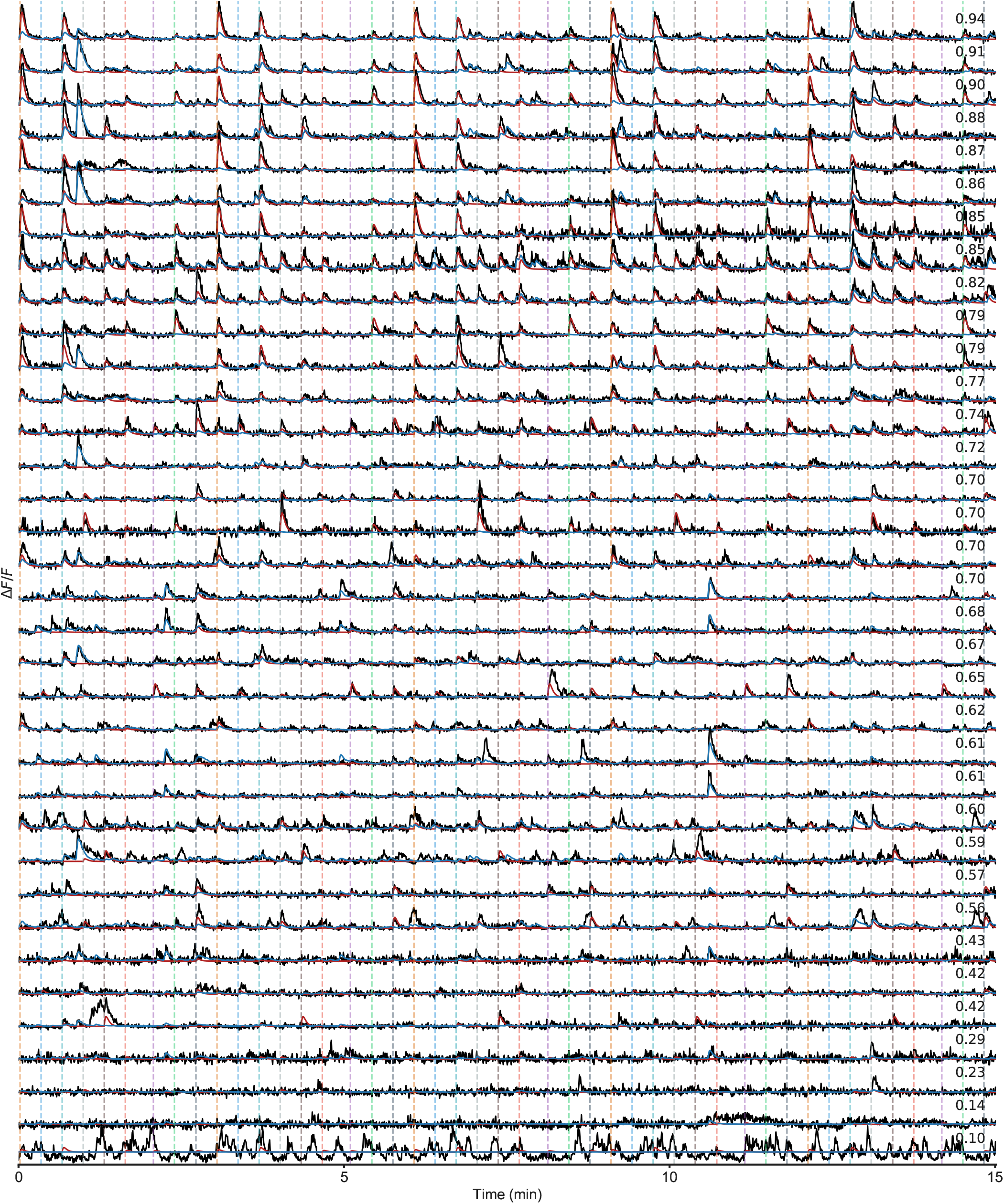
Decoupling of evoked (red) and spontaneous (blue) calcium transients corresponding to the neurons from Supplementary Figure 3.

**Supplementary Figure 5:**
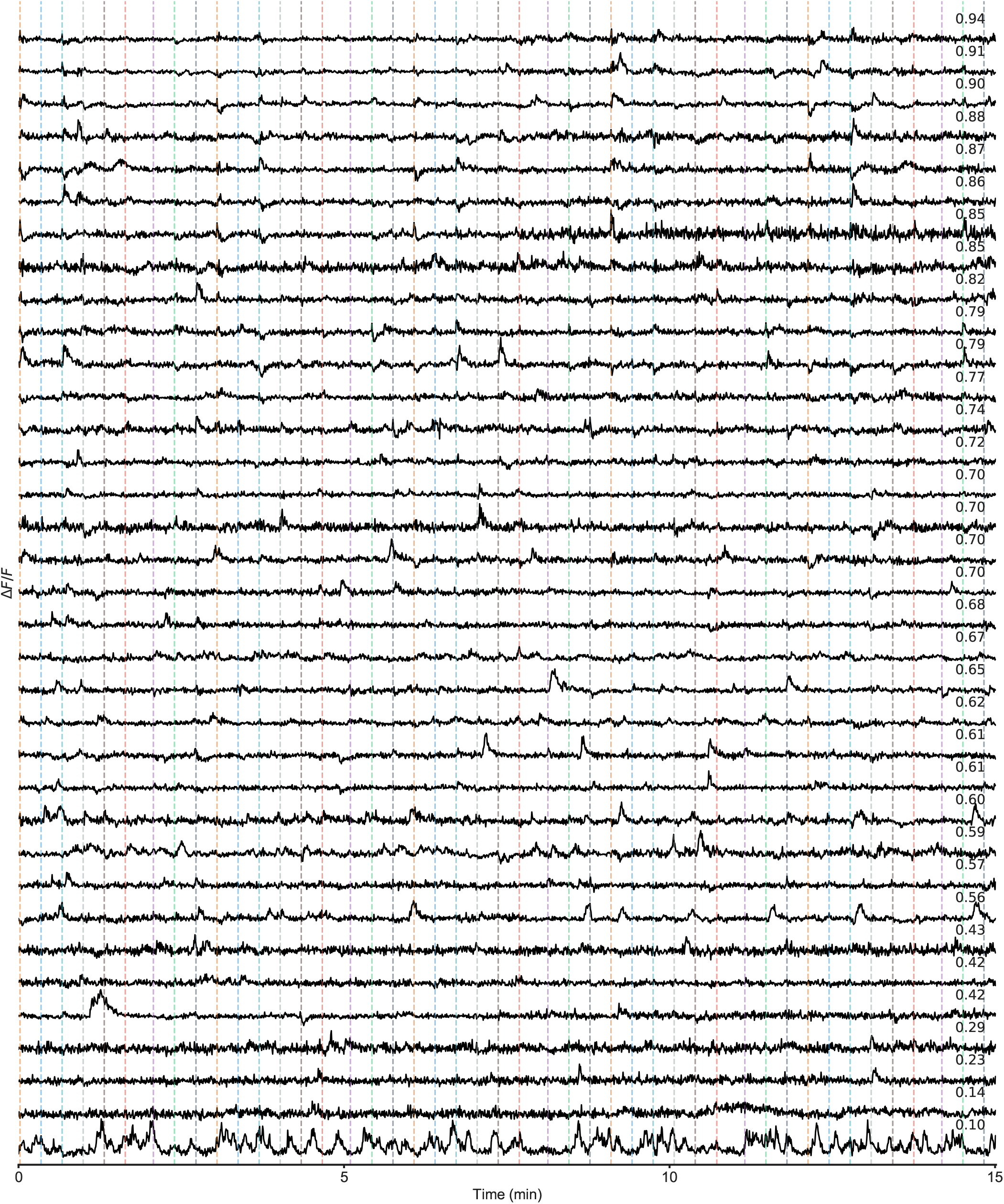
Residuals corresponding to the neurons from Supplementary Figure 3. Residual data obtained by subtracting model fit from the raw data (i.e. 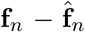). Inset numbers denote the correlation coefficients from the model fits in Supplementary Figure 3. Ideal residuals appear as independent and identically distributed samples from a Gaussian noise distribution. Systematic deviations from Gaussian noise reflect calcium transients not captured by the model, and contribute to measurements of private variability.

**Supplementary Figure 6:**
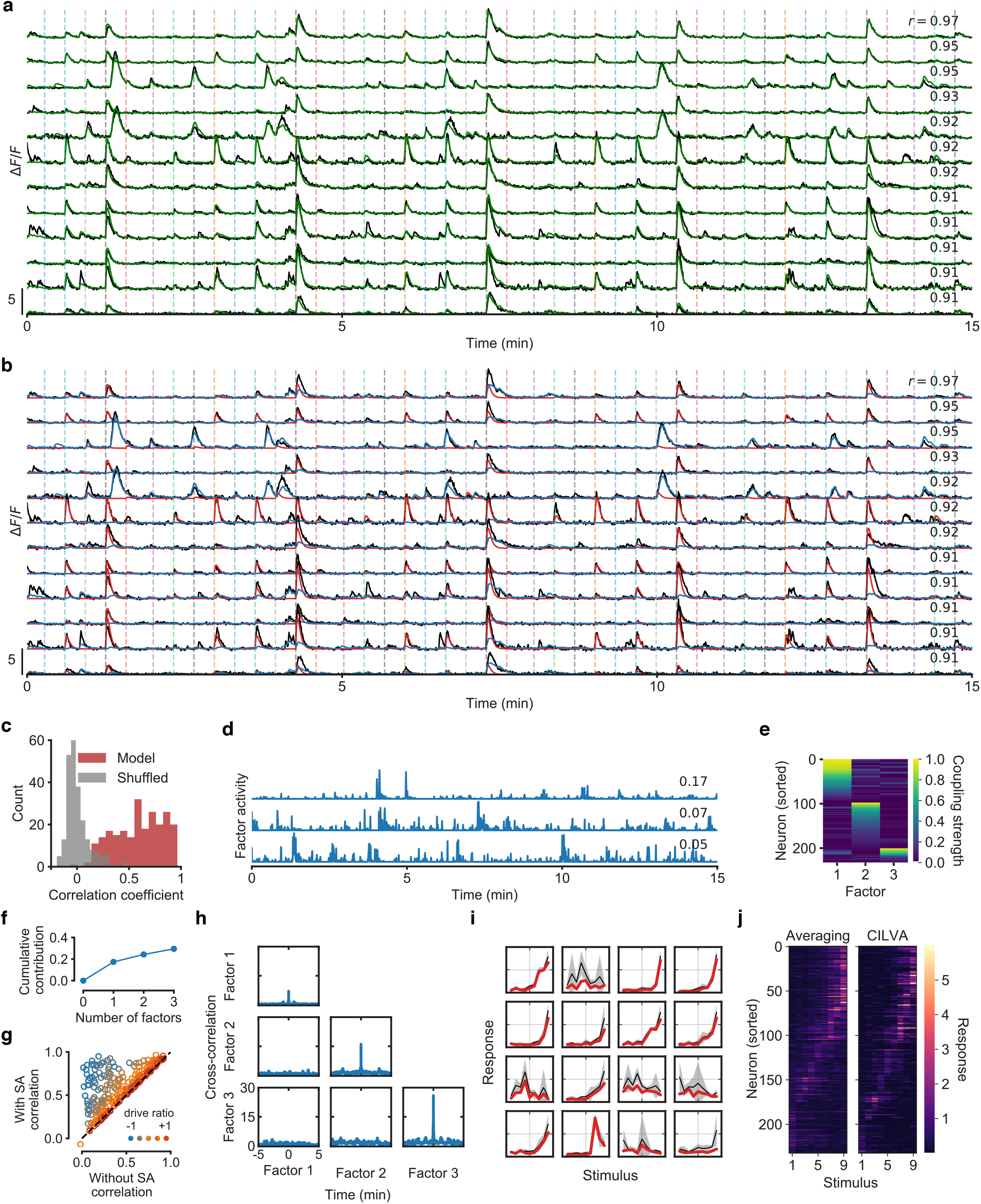
Model fit from a second zebrafish demonstrating similar features to fish shown in the main text. **a**, Example fluorescence traces (black) and model fits (green) for the twelve best fitting neurons. Inset numbers denote the Pearson correlation coefficient between raw trace and model fit. **b**, Application of the statistical model to decouple EA (red) and shared SA (blue). **c**, Distribution of correlation coefficients between data and model fits. Shuffled data (gray) obtained by cyclically permuting each model fit by a random offset while preserving its temporal structure. **d**, Inferred latent factor timeseries. Inset numbers denote the factor contribution indices. **e**, Factor coupling matrix. **f**, Cumulative factor contribution indices for 0-3 latent factors. **g**, Correlation coefficient between raw fluorescence trace and model fit with and without incorporation of SA. Neurons with strongly negative drive ratios show marked improvement in quality of model fit. **h**, Cross-correlograms show little interaction between latent factors. **i**, Example stimulus filters (red). Tuning curves obtained by averaging fluorescence levels over a small window following stimulus presentation provided for comparison (gray). Shaded error bars represent one standard deviation. **j**, Retinotopic maps obtained by averaging (left) and by fitting CILVA (right).

**Supplementary Figure 7:**
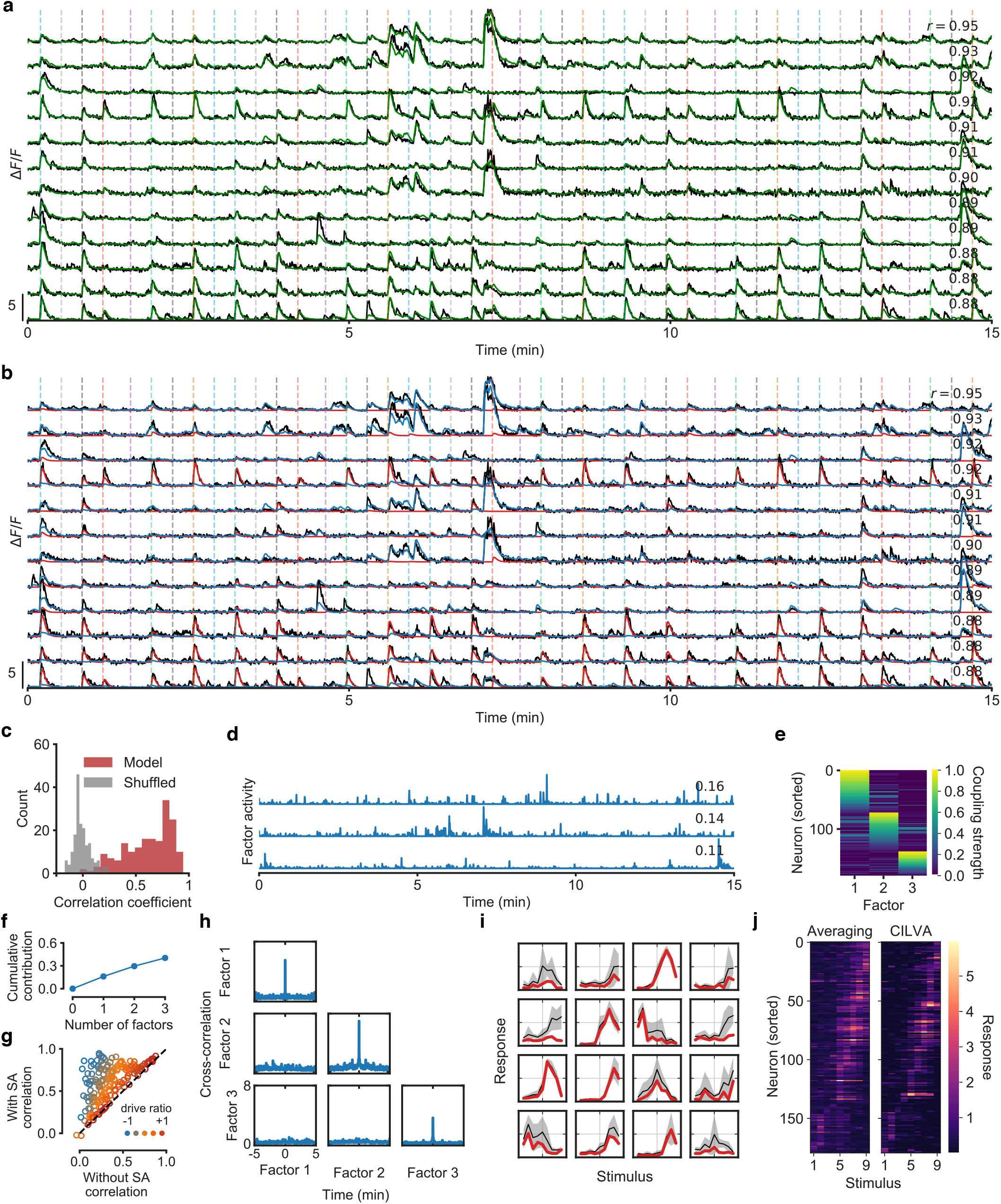
Model fit from a third zebrafish. **a**, Example fluorescence traces (black) and model fits (green) for the twelve best fitting neurons. Inset numbers denote the Pearson correlation coefficient between raw trace and model fit. **b**, Application of the statistical model to decouple EA (red) and shared SA (blue). **c**, Distribution of correlation coefficients between data and model fits. Shuffled data (gray) obtained by cyclically permuting each model fit by a random offset while preserving its temporal structure. **d**, Inferred latent factor timeseries. Inset numbers denote the factor contribution indices. **e**, Factor coupling matrix. **f**, Cumulative factor contribution indices for 0-3 latent factors. **g**, Correlation coefficient between raw fluorescence trace and model fit with and without incorporation of SA. Neurons with strongly negative drive ratios show marked improvement in quality of model fit. **h**, Cross-correlograms show little interaction between latent factors. **i**, Example stimulus filters (red). Tuning curves obtained by averaging fluorescence levels over a small window following stimulus presentation provided for comparison (gray). Shaded error bars represent one standard deviation. **j**, Retinotopic maps obtained by averaging (left) and by fitting CILVA (right).

**Supplementary Figure 8:**
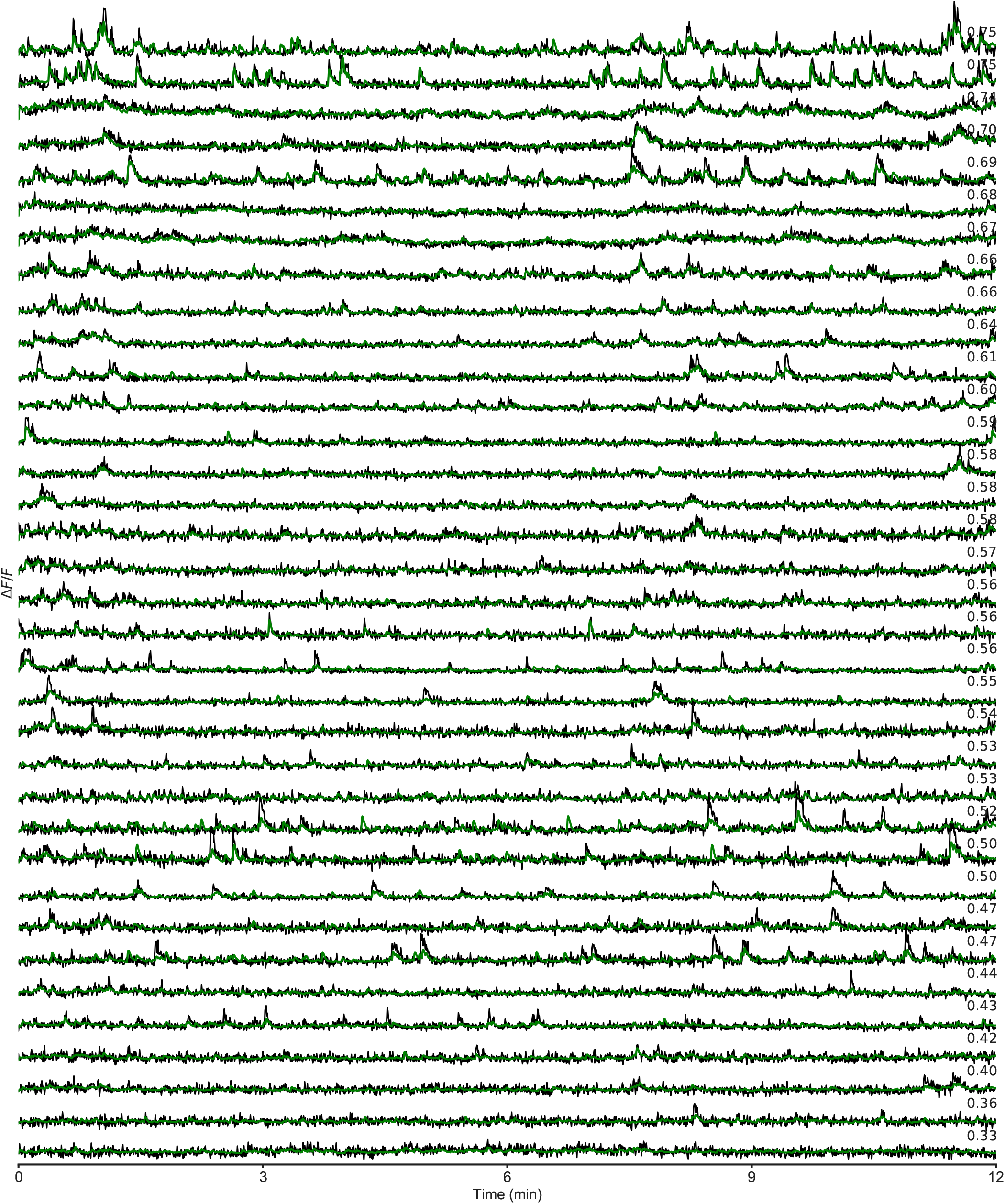
Model fits sampled uniformly from the population of V1 neurons. Example fluorescence traces (black) and corresponding model fits (green). Inset numbers denote Pearson correlation coefficient between raw trace and model fit.

**Supplementary Figure 9:**
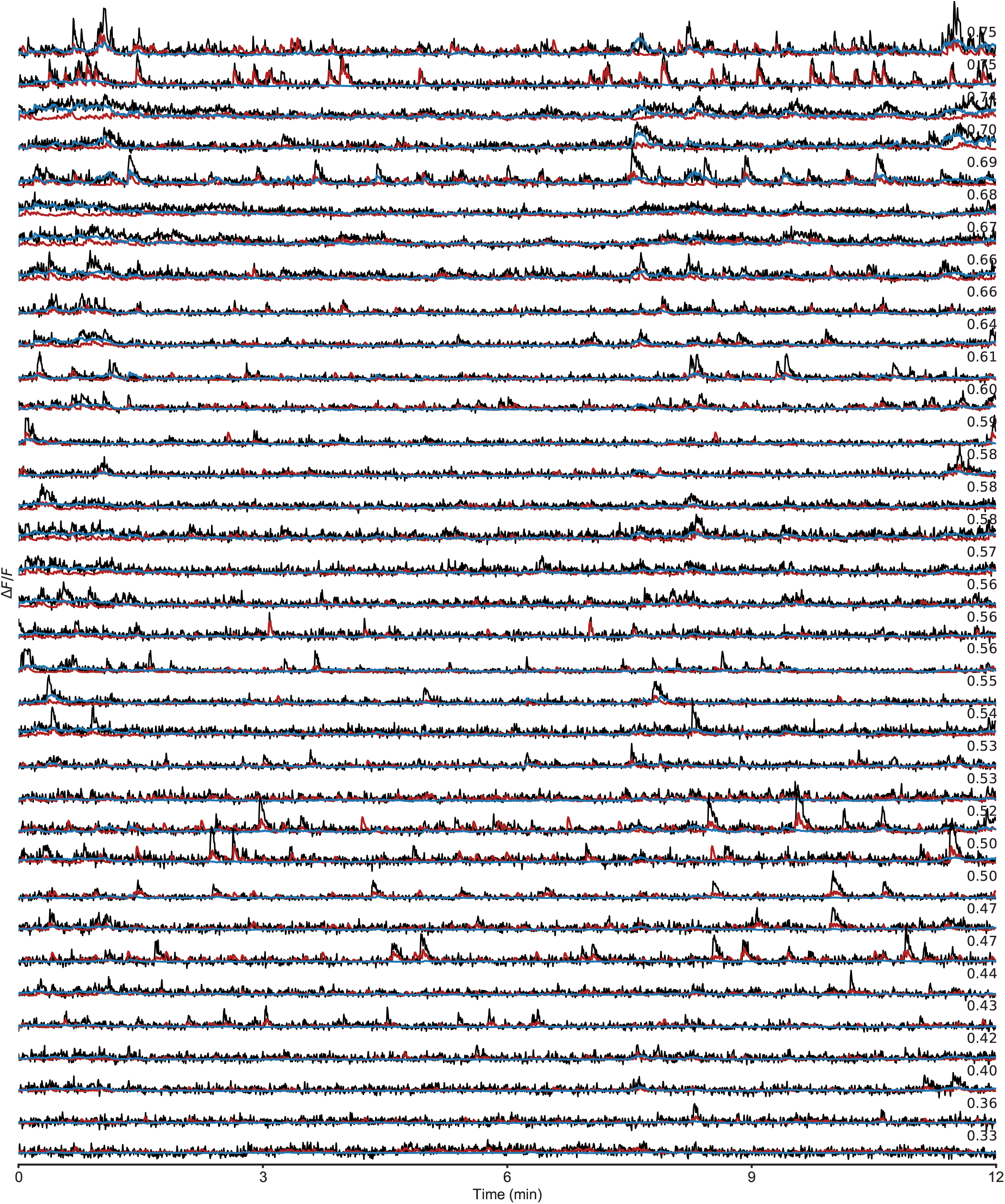
Decoupling of evoked (red) and spontaneous (blue) calcium transients corresponding to the neurons from Supplementary Figure 8.

**Supplementary Figure 10:**
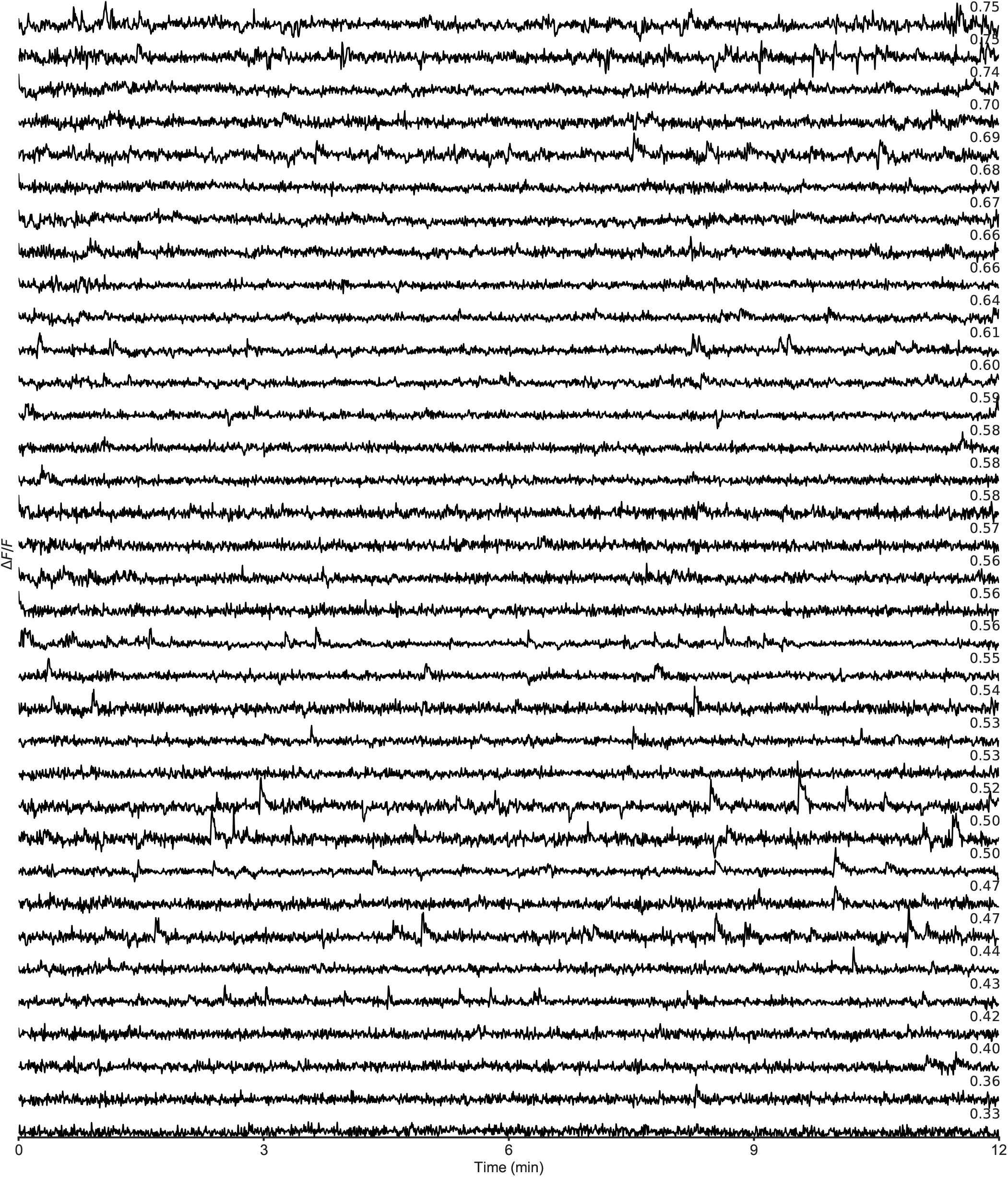
Residuals corresponding to the neurons from Supplementary Figure 8. Residual data obtained by subtracting model fit from the raw data (i.e. 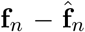). Inset numbers denote the correlation coefficients from the model fits in Supplementary Figure 8.

## References

[1] A. Arieli, A. Sterkin, A. Grinvald, and A. Aertsen. Dynamics of ongoing activity: explanation of the large variability in evoked cortical responses. Science 273.5283 (1996), pp. 1868–1871.

[2] C. Stringer, M. Pachitariu, N. Steinmetz, C. B. Reddy, M. Carandini, and K. D. Harris. Spontaneous behaviors drive multidimensional, brainwide activity. Science 364.6437 (2019).

[3] S. Musall, M. T. Kaufman, A. L. Juavinett, S. Gluf, and A. K. Churchland. Single-trial neural dynamics are dominated by richly varied movements. bioRxiv (2019), p. 308288.

[4] J. B. Ackman, T. J. Burbridge, and M. C. Crair. Retinal waves coordinate patterned activity throughout the developing visual system. Nature 490.7419 (2012), pp. 219–225.

[5] W. E. Allen, M. Z. Chen, N. Pichamoorthy, R. H. Tien, M. Pachitariu, L. Luo, and K. Deisseroth. Thirst regulates motivated behavior through modulation of brainwide neural population dynamics. Science 364.6437 (2019), pp. 253–253.

[6] T.-W. Chen, T. J. Wardill, Y. Sun, S. R. Pulver, S. L. Renninger, A. Baohan, E. R. Schreiter, R. A. Kerr, M. B. Orger, V. Jayaraman, L. L. Looger, K. Svoboda, and D. S. Kim. Ultrasensitive fluorescent proteins for imaging neuronal activity. Nature 499.7458 (2013), pp. 295–300.

[7] J. T. Vogelstein, B. O. Watson, A. M. Packer, R. Yuste, B. Jedynak, and L. Paninski. Spike inference from calcium imaging using sequential Monte Carlo methods. Biophysical Journal 97.2 (2009), pp. 636–655.

[8] L. Paninski and J. P. Cunningham. Neural data science: accelerating the experiment-analysis-theory cycle in large-scale neuroscience. Current Opinion in Neurobiology 50 (2018), pp. 232–241.

[9] M. Y. Byron, J. P. Cunningham, G. Santhanam, S. I. Ryu, K. V. Shenoy, and M. Sahani. Gaussian-process factor analysis for low-dimensional single-trial analysis of neural population activity. Advances in Neural Information Processing Systems. 2009, pp. 1881–1888.

[10] J. H. Macke, L. Buesing, J. P. Cunningham, M. Y. Byron, K. V. Shenoy, and M. Sahani. Empirical models of spiking in neural populations. Advances in Neural Information Processing Systems. 2011, pp.1350–1358.

[11] J. P. Cunningham and M. Y. Byron. Dimensionality reduction for large-scale neural recordings. Nature Neuroscience 17.11 (2014), pp. 1500–1509.

[12] C. Pandarinath, D. J. O’Shea, J. Collins, R. Jozefowicz, S. D. Stavisky, J. C. Kao, E. M. Trautmann, M. T. Kaufman, S. I. Ryu, L. R. Hochberg, et al. Inferring single-trial neural population dynamics using sequential auto-encoders. Nature Methods 15 (2018), pp. 805–815.

[13] P. T. Sadtler, K. M. Quick, M. D. Golub, S. M. Chase, S. I. Ryu, E. C. Tyler-Kabara, M. Y. Byron, and A. P. Batista. Neural constraints on learning. Nature 512.7515 (2014), pp. 423–426.

[14] L. Aitchison, L. Russell, A. M. Packer, J. Yan, P. Castonguay, M. Häusser, and S. C. Turaga. Model-based Bayesian inference of neural activity and connectivity from all-optical interrogation of a neural circuit. Advances in Neural Information Processing Systems. 2017, pp. 3489–3498.

[15] E. Kirschbaum, M. Haußmann, S. Wolf, H. Sonntag, J. Schneider, S. Elzoheiry, O. Kann, D. Durstewitz, and F. A. Hamprecht. LeMoNADe: Learned Motif and Neuronal Assembly Detection in calcium imaging videos. International Conference on Learning Representations. 2019.

[16] A. Wu, S. Pashkovski, S. R. Datta, and J. W. Pillow. Learning a latent manifold of odor representations from neural responses in piriform cortex. Advances in Neural Information Processing Systems 31. 2018, pp. 5378–5388.

[17] T. O. Helmbrecht, M. Dal Maschio, J. C. Donovan, S. Koutsouli, and H. Baier. Topography of a Visuomotor Transformation. Neuron 100.6 (2018), pp. 1429–1445.

[18] X. Chen, Y. Mu, Y. Hu, A. T. Kuan, M. Nikitchenko, O. Randlett, A. B. Chen, J. P. Gavornik, H. Sompolinsky, F. Engert, et al. Brain-wide organization of neuronal activity and convergent sensorimotor transformations in larval zebrafish. Neuron 100.4 (2018), pp. 876–890.

[19] D. D. Lee and H. S. Seung. Learning the parts of objects by non-negative matrix factorization. Nature 401.6755 (1999), pp. 788–791.

[20] E. A. Pnevmatikakis, D. Soudry, Y. Gao, T. A. Machado, J. Merel, D. Pfau, T. Reardon, Y. Mu, C. Lacefield, W. Yang, et al. Simultaneous Denoising, Deconvolution, and Demixing of Calcium Imaging Data. Neuron 89.2 (2016), pp. 285–299.

[21] G. Santhanam, B. M. Yu, V. Gilja, S. I. Ryu, A. Afshar, M. Sahani, and K. V. Shenoy. Factor-analysis methods for higher-performance neural prostheses. Journal of Neurophysiology 102.2 (2009), pp. 1315–1330.

[22] M. R. Whiteway and D. A. Butts. Revealing unobserved factors underlying cortical activity with a rectified latent variable model applied to neural population recordings. Journal of Neurophysiology 117.3 (2016), pp. 919–936.

[23] I.-C. Lin, M. Okun, M. Carandini, and K. D. Harris. The nature of shared cortical variability. Neuron 87.3 (2015), pp. 644–656.

[24] A. Litwin-Kumar and B. Doiron. Slow dynamics and high variability in balanced cortical networks with clustered connections. Nature Neuroscience 15.11 (2012), pp. 1498–1505.

[25] M. A. Triplett, L. Avitan, and G. J. Goodhill. Emergence of spontaneous assembly activity in developing neural networks without afferent input. PLoS Computational Biology 14.9 (2018), e1006421.

[26] A. W. Thompson, G. C. Vanwalleghem, L. A. Heap, and E. K. Scott. Functional profiles of visual-, auditory-, and water flow-responsive neurons in the zebrafish tectum. Current Biology 26.6 (2016), pp. 743–754.

[27] M. Pachitariu, C. Stringer, and K. D. Harris. Recordings of 10k neurons in V1 during drifting gratings. May 2018. DOI: 10.25378/janelia.6214019.v1.

[28] K. V. Shenoy, M. Sahani, and M. M. Churchland. Cortical control of arm movements: a dynamical systems perspective. Annual Review of Neuroscience 36 (2013), pp. 337–359.

[29] D. Shimaoka, N. A. Steinmetz, K. D. Harris, and M. Carandini. The impact of bilateral ongoing activity on evoked responses in mouse cortex. eLife 8 (2019), e43533.

[30] M. Vidne, Y. Ahmadian, J. Shlens, J. W. Pillow, J. Kulkarni, A. M. Litke, E. Chichilnisky, E. Simoncelli, and L. Paninski. Modeling the impact of common noise inputs on the network activity of retinal ganglion cells. Journal of Computational Neuroscience 33.1 (2012), pp. 97–121.

[31] M. R. Deweese and A. M. Zador. Shared and private variability in the auditory cortex. Journal of Neurophysiology 92.3 (2004), pp. 1840–1855.

[32] M. Okun, N. A. Steinmetz, L. Cossell, M. F. Iacaruso, H. Ko, P. Barthó, T. Moore, S. B. Hofer, T. D. Mrsic-Flogel, M. Carandini, et al. Diverse coupling of neurons to populations in sensory cortex. Nature 521.7553 (2015), pp. 511–515.

[33] E. K. Miller and J. D. Cohen. An integrative theory of prefrontal cortex function. Annual Review of Neuroscience 24.1 (2001), pp. 167–202.

[34] L. Petreanu, D. A. Gutnisky, D. Huber, N.-l. Xu, D. H. O’connor, L. Tian, L. Looger, and K. Svoboda. Activity in motor–sensory projections reveals distributed coding in somatosensation. Nature 489.7415 (2012), pp. 299–303.

[35] J. Gründemann, Y. Bitterman, T. Lu, S. Krabbe, B. F. Grewe, M. J. Schnitzer, and A. Lüthi. Amygdala ensembles encode behavioral states. Science 364.6437 (2019), eaav8736.

[36] M. Westerfield. The Zebrafish Book: A Guide for the Laboratory Use of Zebrafish (Brachydanio rerio). University of Oregon Press, 2000.

[37] M. Dipoppa, A. Ranson, M. Krumin, M. Pachitariu, M. Carandini, and K. D. Harris. Vision and locomotion shape the interactions between neuron types in mouse visual cortex. Neuron 98.3 (2018), pp. 602–615.

[38] M. Pachitariu, C. Stringer, and K. D. Harris. Robustness of spike deconvolution for neuronal calcium imaging. Journal of Neuroscience 38.37 (2018), pp. 7976–7985.

[39] F. Pedregosa, G. Varoquaux, A. Gramfort, V. Michel, B. Thirion, O. Grisel, M. Blondel, P. Prettenhofer, R. Weiss, V. Dubourg, et al. Scikit-learn: Machine learning in Python. Journal of Machine Learning Research 12.Oct (2011), pp. 2825–2830.

[40] J. T. Vogelstein, A. M. Packer, T. A. Machado, T. Sippy, B. Babadi, R. Yuste, and L. Paninski. Fast nonnegative deconvolution for spike train inference from population calcium imaging. Journal of Neurophysiology 104.6 (2010), pp. 3691–3704.

